# Visual Semantic Encoding and Identification of Naturalistic Movies via High-Density Diffuse Optical Tomography

**DOI:** 10.64898/2025.12.03.692158

**Authors:** Wiete Fehner, Morgan Fogarty, Jerry Tang, Dana Wilhelm, Aahana Bajracharya, Zachary E. Markow, Amelia Hines, Jason W. Trobaugh, Alexander G. Huth, Joseph P. Culver

## Abstract

Understanding how the brain represents meaning in real-world contexts is essential for both fundamental neuroscience and clinical applications. Brain encoding and decoding models from naturalistic stimuli provide a powerful window into semantic representations. Yet, existing approaches rely on a constrained scanning environment, or on conventional fNIRS, which has been limited to sparse sampling and/or block-design paradigms. Here, we tested whether high-density diffuse optical tomography (HD-DOT), an advanced high-density tomographic optical imaging method, can support semantic encoding and decoding using naturalistic movies. We collected 3.5 hours of naturalistic movie viewing data from six participants using stimuli labeled with 1,708 categories. Encoding models robustly predicted voxel-level responses, yielding single semantic category maps consistent with prior fMRI studies. In complementary decoding analyses, we showed that DOT responses captured sufficient semantic content to identify which clips participants viewed. To assess organization across individuals, we identified a shared low-dimensional semantic space that captures common semantic dimensions. Finally, clustering analyses revealed interpretable higher-order semantic dimensions like social and animate agents, objects vs natural organisms, and textural scenes, consistently mapped across the cortex. These findings demonstrate that DOT can recover distributed, high-dimensional semantic representations from naturalistic movies, bridging fMRI-level semantic mapping with the accessibility of optical imaging.

## Introduction

Understanding meaning in the real world involves extracting semantic information, such as objects, actions, and social cues, from continuous visual scenes. This process of deriving meaning from visual experience, or visual semantics, has been extensively studied using many neuroscience techniques [1–7]. Recent work has shown that naturalistic paradigms capture perception and meaning more accurately as they unfold continuously over time in real-world contexts, rather than in isolated, static events [8, 9]. Building on this shift, a rich set of fMRI studies has pioneered naturalistic semantic mapping, demonstrating that distributed cortical responses encode high-dimensional semantic features during movie viewing and podcast listening [10–17]. These advances rest on decades of foundational work in movie-based paradigms [18–21], detailed language mapping [22, 23], and visual paradigms [24–27], as well as collecting many hours of data from single participants [28–30]. Despite these breakthroughs, the constrained imaging environment of MRI limits ecological, naturalistic, and clinical applications [31–33].

Optical neuroimaging techniques, including functional near-infrared spectroscopy (fNIRS) and its tomographic extension, Diffuse Optical Tomography (DOT), have become a promising and powerful alternative to fMRI while providing a more open scan environment and enabling use in more naturalistic settings [34–42]. The vast majority of the fNIRS literature has relied on sparse imaging arrays (<50 channels) that lack the image quality necessary for advanced semantic mapping. But recent fNIRS and DOT studies have been building experience and tools that point towards high-dimensional semantic experiments. Early fNIRS studies with sparse arrays and block-design paradigms demonstrated that semantic information can be encoded and decoded, using coarse semantic object-category contrasts [43–49]. High-Density Diffuse Optical Tomography (HD-DOT) systems that employ imaging arrays (>1000 channels) have demonstrated functional mapping with a fidelity much closer to that of fMRI [34–36]. And recently, a collection of HD-DOT studies has assembled the enabling components for advanced semantic mapping, including visual paradigms [36, 50, 51], language-mapping approaches [35, 52, 53], movie-mapping frameworks [40, 51, 54–56], and precision multi-session imaging methods [57]. While progress has been made, movie-based HD-DOT studies have remained limited to short movies (∼20 min) and to only a few semantic categories.

Here, we developed a high-dimensional semantic encoding and decoding framework for HD-DOT. We collected 3.5 hours of naturalistic movies per subject and employed rich annotation with 1,708 semantic categories. Using this dataset, we mapped visual semantic representations at multiple levels of organization, including: (1) voxelwise encoding with high-dimensional features to model large-scale semantic representation, (2) single category cortical maps to reveal spatial patterns of semantic representations, and (3) a shared low-dimensional space across participants that captures distributed, higher-order semantic structure. To assess the semantic information content in the DOT data, we evaluated model-based decoding of clip identification. To evaluate whether the semantic maps were consistent with principles of cortical organization, we applied dimensionality reduction and clustering, which enabled visualization of the semantic content and its interpretability. By adapting high-dimensional semantic modeling to DOT, we extend naturalistic semantic neuroscience beyond the MRI scanner, opening new opportunities in real-world settings and for populations difficult to access with fMRI.

## Results

### Diffuse Optical Tomography

Participants were scanned using a recently developed very-high-density DOT system, with ∼9.75 mm spacing between first-nearest-neighbor source and detector pairs (SD-pairs) [40]. The cap consists of a total of 255 sources and 252 detectors. Using SD-pairs with distances ≤40 mm, there are 9,160 possible SD-pair measurements. Retained measurements satisfied the temporal variance threshold (<7.5%) applied to each run (**Figure S1E**, **Figure S1G**, **Figure S1H**). Across participants, the number of retained SD-pair measurements ranged from 6,917 to 8,598 (86.7% ± 2.6%, mean ± SEM measurements retained).

Each SD-pair measurement reflects average light attenuation through a diffuse sampling of the head. Subject-specific light models (for the full set of sources and detectors) were constructed using a combination of MRI anatomy, photogrammetric data, and functional localizers. Each subject’s model was inverted to transform SD-pair measurements into three-dimensional voxelwise reconstructions. These tomographic images provide voxelwise differential absorption across the scalp, skull, and cortical tissue [58]. Overlap field-of-view (FOV) maps, obtained from subject-specific light modeling (see **Methods**), confirmed consistent whole-head coverage for all participants (**Figure S1B, Figure S1F**).

### Semantic Encoding Model Validation

To establish a semantic encoding framework with DOT, we had six participants view naturalistic movies. Each participant viewed 210 minutes (3.5 hours) of movies, including 120 minutes of unique movies for training, and 9 minutes of movies that were repeated ten times for model testing, allowing robust training, validation, and evaluation (**Figure S1A**, **Figure S1C**). This paradigm and stimulus set match those used in earlier fMRI studies [12, 14, 21]. To make this paradigm more comfortable for participants, we spread the imaging across three same-day sessions, using functional localizers within each session to provide alignment measures. To assess cross-session repeatability, we quantified correlations of visual and auditory localizer responses across sessions and evaluated explained variance for test movie repeats. These analyses demonstrated robust repeatability for both localizers and movie responses in four participants, with two participants showing lower repeatability (**Figure S2A-C**). With the aim of capturing visual semantics, each second of the movie stimulus was manually labeled with nouns and verbs describing its contents, giving a total of 1,708 action and object categories following the Wordnet taxonomy [59]. To link semantic features with brain activity, we used L2-regularized linear regression to estimate a set of encoding model coefficients for each voxel. The encoding model coefficients predict how the presence of each semantic category influences DOT time courses (**Figure 1A**). To account for hemodynamic delays, we used a finite impulse response (FIR) basis, where the original feature-by-time matrix included four delays capturing 1 to 8 seconds (**Figure 1A-B**).

**Figure 1.**
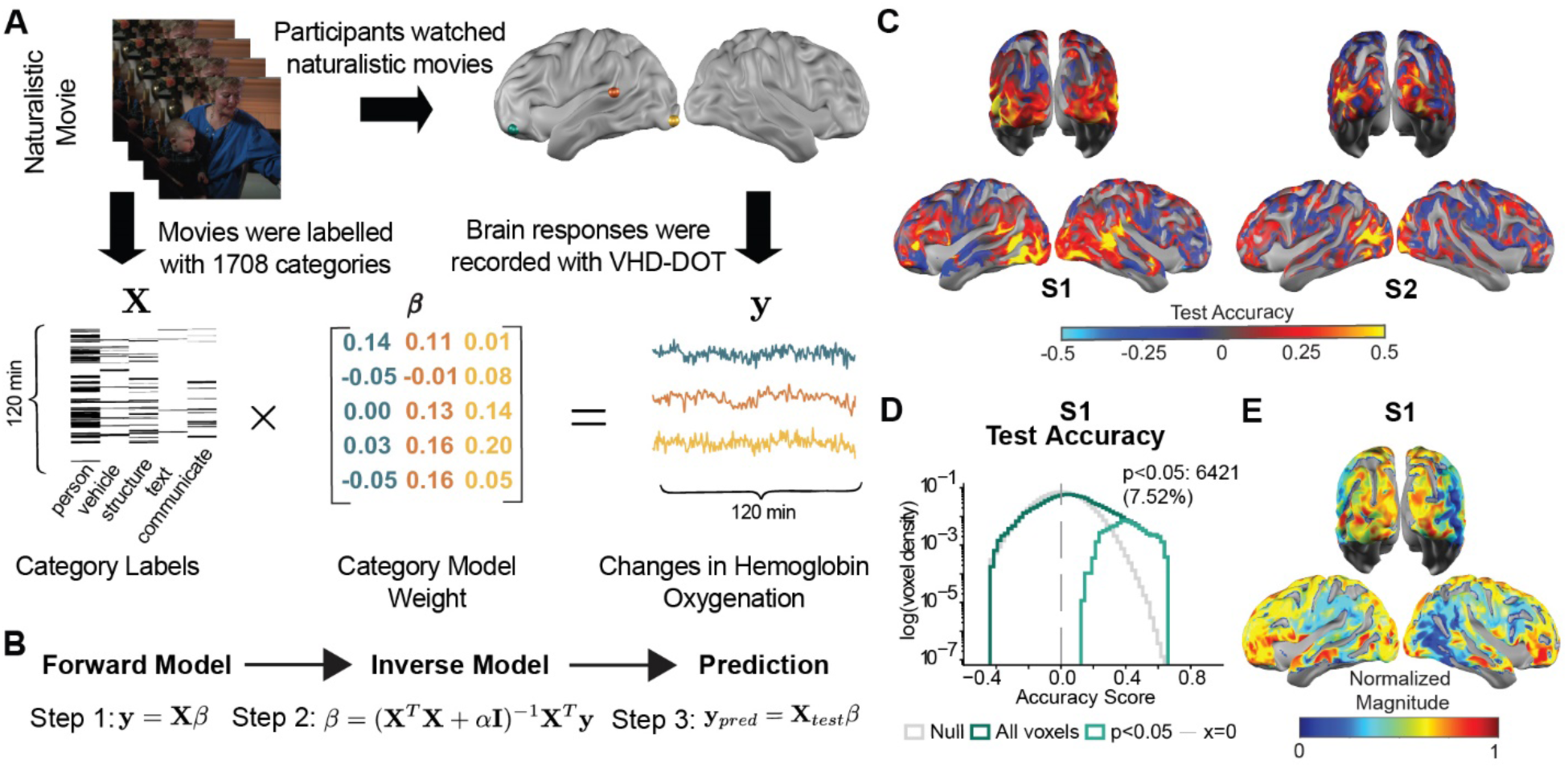
| Semantic encoding paradigm, model analysis and validation. **(A)** The semantic encoding model predicts voxel-level brain activity from the semantic features of a naturalistic movie stimulus. **(B)** Model weights (𝜷) were estimated using L2-regularized linear regression with a finite impulse response (FIR) model that included four time delays to capture the hemodynamic response. **(C)** Voxelwise model accuracy maps (Pearson correlation between predicted, 𝒚_𝒑𝒓𝒆𝒅_, and actual responses, 𝒚) are shown for two subjects, evaluated on held-out test data. **(D)** Test accuracy was determined through correlation of the predicted and observed DOT response. Voxelwise significance was determined using a 1,000-iteration blockwise permutation test (60 s blocks) with FDR correction (p < 0.05). Correlation values were interpreted relative to the null distribution, where higher positive correlations indicate better model fit than expected by chance. Also, see **Figure S2D** for the remaining subjects. **(E)** Spatial distribution of normalized weight magnitude (RMS) for one subject. Weights were first averaged across FIR delays and then normalized per voxel to focus on semantic selectivity pattern rather than magnitude. The resulting map reveals strong weights in occipital and frontal regions, consistent with higher-order semantic networks. See **Figure S1** for the experimental design, imaging system and data quality.

To evaluate the models, test accuracy was quantified on held-out test data using Pearson correlations between predicted and observed time courses (**Figure 1C**). Voxelwise significance was determined using a 1,000-iteration blockwise permutation test (60 s blocks) with FDR correction (p < 0.05). Correlation values were interpreted relative to the null distribution, where higher positive correlations indicate better model fit than expected by chance. Distributions of voxelwise test accuracy, corresponding null distributions, and significant voxels are shown for all subjects (**Figure 1D, Figure S2D**). Across participants, mean test accuracy was 0.032 ± 0.008 compared to a null mean of 0.004 ± 0.001 (mean ± SEM). Four participants exhibited significant voxels (2471 ± 1154, corresponding to 3.12% ± 1.31% of voxels), while two participants showed symmetric distributions of test accuracy without significant voxels. Nonetheless, semantic information in the DOT signal is clearly evident in all participants, as demonstrated by above-chance clip identification decoding performance (**Figures 3**, **Figure 4**). To illustrate cortical localization of model performance, voxelwise test accuracy maps from two participants are shown, exhibiting widespread regions of high accuracy, including lateral occipitotemporal and frontal cortex (**Figure 1C**).

To obtain interpretable model coefficients, we averaged across time delays, yielding a single weight for each voxel and each category. A single regularization parameter (𝜶) was used for all voxels, allowing direct comparison of coefficient magnitudes across the cortex. To characterize cortical locations of semantic sensitivity, we first visualized the normalized root mean square (RMS) magnitude of model weights for each voxel. Strong weightings indicate larger sensitivity to the semantic features. Strongest weightings emerged in occipital regions associated with visual processing and in frontal regions implicated in higher-order semantic networks (**Figure 1E**). This pattern is consistent with distributed visual-semantic representations observed in prior fMRI studies [12, 16].

### Single Semantic Category Mapping

To test whether individual semantic categories could be resolved, we mapped voxel-level model weights at the group level (**Figure 2**) and individual level (**Figure S3**). For the group-level analysis, we used fixed-effect t-statistics computed across participants from the encoding model weights (averaged over delays) for five categories: *person*, *communicate*, *text*, *structure*, and *vehicle* (**Figure 2A-E**). We focused on these five representative categories because they cover diverse semantic domains (social, linguistic, symbolic, environmental, and object-related) and engage distinct cortical networks [5, 10, 12, 16, 17].

**Figure 2.**
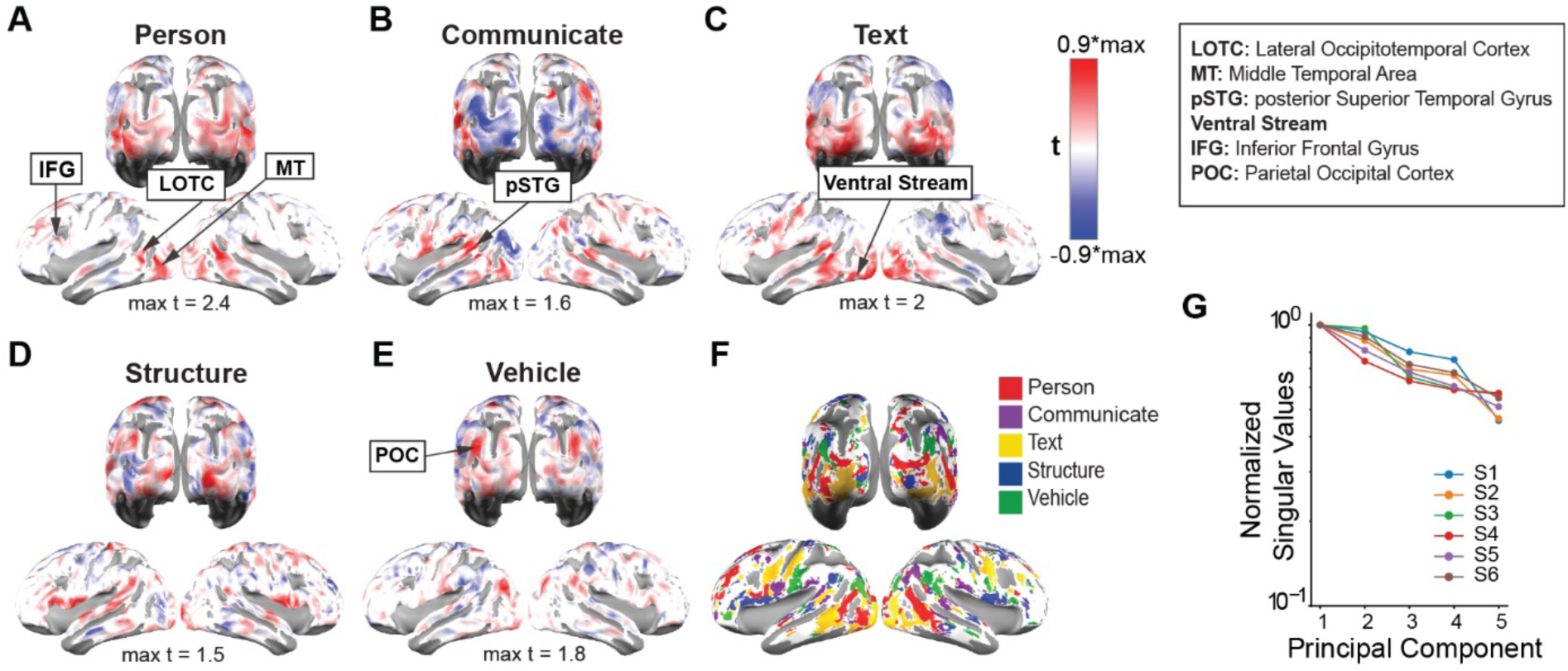
| Single Semantic Category Mapping. (A-E) Group T-maps (fixed effect) for five semantic categories mapped onto MNI space. **(F)** A winner-takes-all strategy was applied to identify the dominant category at each voxel and map all five categories simultaneously. Voxels with low t-value magnitude (𝒕 < 0.2 ∗ 𝑚𝑎𝑥|𝒕| or ambiguous category preference (i.e., small difference (< 0.1) between the top two categories) were rendered white. Each voxel was assigned a distinct RGB color based on the winning category. The resulting map was projected onto the cortical surface for visualization. **(G)** PCA was performed on a 5 × n voxels matrix of the five category weights for each subject. The plot shows normalized singular values for six subjects, indicating that variance is spread across multiple components, with distinct spatial patterns for the five categories.

**Figure 3.**
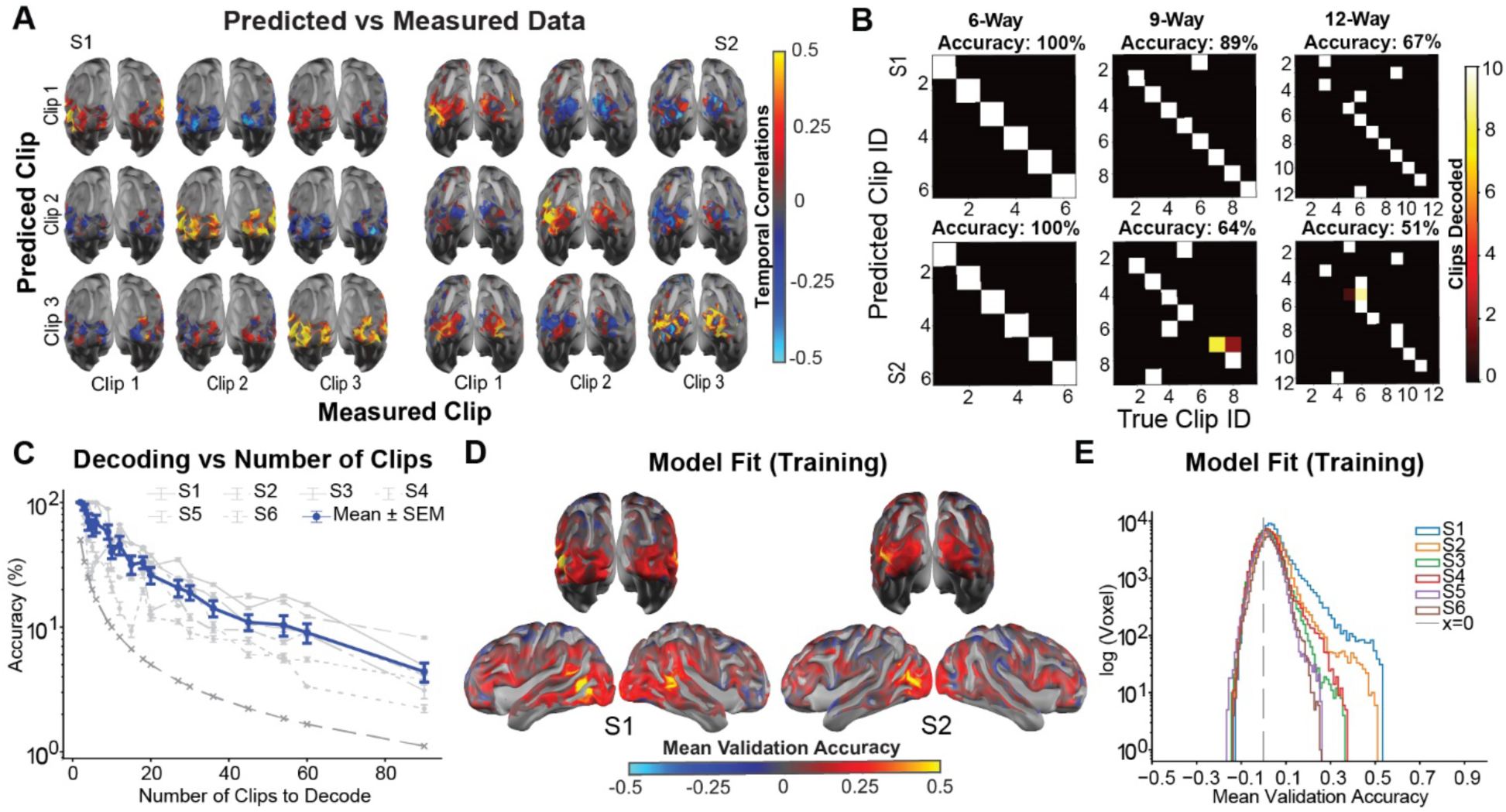
| Model-based clip decoding with semantic features. **(A)** Voxelwise temporal correlation for two example subjects, showing higher correlations for matched than mismatched conditions across three 3-min test clips between predicted and measured responses. Decoding was performed on the top 10% most reliable voxels, identified by a bootstrap procedure on training data. See **Figure S5** for the lateral view and the remaining subjects. **(B)** Clip decoding at finer segmentations (90 s, 60 s, 30 s) for two subjects aggregated over bootstrapping repetitions. Above-chance diagonal structure is consistent across all granularities. See **Figure S6** for all subjects. **(C)** Group decoding accuracy as a function of the number of clips to identify. Nine minutes of test data were segmented into 2 to 90 clips (270 to 6 s). Robust decoding was maintained across granularities. **(D)** Training accuracy maps for two subjects were used to define the voxel mask. **(E)** Group distribution of training accuracy across the top 10% selected voxels, showing consistently positive correlations across subjects. See **Methods** for voxel selection procedure.

**Figure 4.**
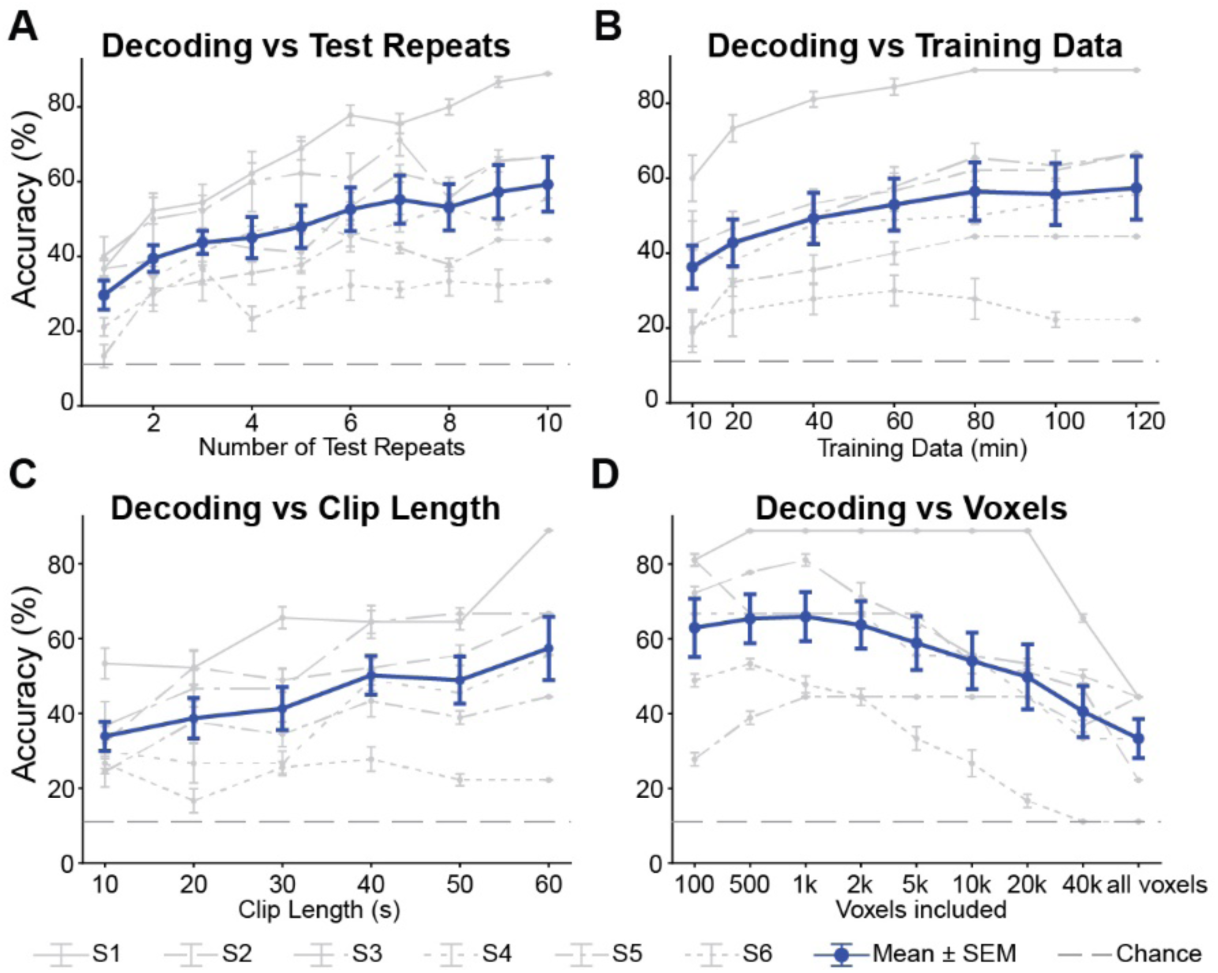
| Clip Identification Decoder Robustness (9-way decoding task). **(A)** Decoding accuracy improves as more repeated presentations are averaged in the test set. Models were fixed, only the test data varied. **(B)** Training data was systematically reduced to 10, 20, 40, 60, 80, and 100 minutes (1 to 12 runs of 10 min each), with run order randomized and repeated 10 times to preserve temporal structure. Encoding models were fit for each fraction, and a clip identification task was performed. Decoding accuracy was evaluated on held-out test data and averaged across bootstrap repetitions. **(C)** Accuracy increases with test clip length (10 to 60 s), reflecting greater discriminability. **(D)** Voxel selection thresholding shows that decoding is driven by the top-performing voxels. All voxel selection was based on bootstrapped training accuracy to avoid test data leakage. All panels show single subject performance (gray lines) and group mean ± SEM (blue lines). For single subject performance, mean ± SEM was computed across bootstrap iterations.

Also, see **Figure S3** for single-subject category maps and **Figure S4** for reproducibility analysis of single categories on a single subject and group level.

For *person*, high model weights emerged in regions associated with social and body perception, including the inferior frontal gyrus (IFG), the lateral occipitotemporal cortex (LOTC) [4, 6], and the motion-sensitive middle temporal area (MT) [60] (**Figure 2A**). For *communicate*, activation was localized to language-related regions, including the posterior superior temporal gyrus (pSTG), which is consistent with visual speech perception [61, 62] and social interaction [63] (**Figure 2B**). For *text*, the strongest responses were found in the ventral visual stream, which is implicated in object and word recognition [64] (**Figure 2C**). Responses to the *structure* category were more diffuse, with activations in occipital regions, reflecting scene and object processing [5, 17] (**Figure 2D**). For the *vehicle* category, activation patterns overlapped with the parietal occipital cortex (POC), a region sensitive to object-rich categories [5] (**Figure 2E**). This pattern likely reflects the distributed nature of scene and texture representations, which in fMRI engage multiple regions (occipital place area (OPA), parahippocampal place area (PPA), retrosplenial cortex (RSC)), of which only OPA lies within the HD-DOT field-of-view [5, 65].

To visualize single-subject responses, model weights for three example categories (*person*, *communicate*, *text*) were projected onto cortical surfaces for individual participants. These showed qualitative consistency with group t-maps, highlighting reproducibility of category-selective patterns (**Figure S3**). Winner-takes-all maps of the five categories revealed distinct single-category patterns that were consistent across subjects (**Figure 2F**). To quantitatively test the independence of semantic category representations, PCA was performed on the matrix of category weights (5 × n voxels) for each subject. A slow decay in the singular value spectrum would indicate that components are relatively orthogonal rather than dominated by a shared single dimension. The resulting singular value spectra showed variance distributed across multiple components, consistent with a distinct and non-redundant representation of individual semantic categories (**Figure 2G**).

To assess the reproducibility of category weights, we implemented a category detection analysis using 50:50 splits of the training data. Regression models were fit separately to each split, and category weights were correlated across splits. A category was considered repeatably detected when its cross-split correlation exceeded that of all other categories. This yielded above-chance accuracy for every participant (range: 41.6%-60.0%, mean ± SEM; chance = 20%) and at the group level (54.2% ± 2.9%) (**Figure S4**). These results demonstrate that DOT reliably captures non-redundant, distributed semantic structure at the level of individual categories.

### Model-based Semantic Decoding

While encoding analyses quantify how well semantic features explain DOT responses, decoding assesses how much semantic information is contained in the DOT signal. We evaluated DOT responses by implementing a semantic model-based clip identification decoding approach to determine whether semantic structure could be recovered from brain activity during naturalistic movie viewing (**Figures 3**, **Figure 4**). To constrain decoding to informative voxels, voxelwise training accuracy maps were generated from cross-validation results obtained during model training (**Figure 3E**, **Figure 3D**; see **Methods**). The top 10% of voxels were selected and used to build voxel masks for decoding. Voxel masks were built exclusively from training folds, per subject. To evaluate decoding performance at the single-subject level, we computed temporal correlation between predicted and measured voxel responses for a 3-way decoding task from the 9-minute held-out test data. Voxelwise surface maps revealed high correlation for the predicted and true clips in the top participants (**Figure 3A**, **Figure S5**). To test how decoding performance scales with temporal granularities, we evaluated increasing numbers of clips spanning the same 9 min test set. At intermediate granularities (6-way, 9-way, 12-way), spatiotemporal correlation produced consistent above-chance decoding. Average decoding accuracies were 68% ± 11% (6-way, chance = 17%), 58% ± 8% (9-way, chance = 11%), and 46% ± 8% (12-way, chance = 8%) (mean ± SEM; **Figure 2B, Figure S6**). Finally, to systematically assess decoding performance across a full range of temporal resolutions, we varied the number from 2 clips (270 s each) to 90 clips (6 s each). Accuracy declined with shorter segments, but the performance was consistently higher than chance, indicating that semantic structure remains recoverable even with limited temporal context (**Figure 3C**). Together, these results suggest that decoding is feasible and remains robust across different temporal segmentations.

To further systematically assess the limits of clip identification decoding, we independently varied test data averaging, training data size, clip length, and voxel inclusion for a 9-way decoding task. The group mean decoding accuracy was well above chance across conditions (**Figure 4**). To examine the strength of DOT movie responses relative to spurious DOT signal in decoding, we varied the number of repeated test clips averaged. Accuracy increased with more repetitions (**Figure 4A**). To evaluate the effect of training data size, decoding was performed with subsets of 10 minutes runs (1, 2, 4, 6, 8, 10, 12 of 12 runs). Performance increased on average with additional training data but plateaued around 80 minutes (56.5% ± 7% accuracy; chance = 11%, mean ± SEM). This saturation was driven largely by near-ceiling performance in the top subjects (**Figure 4B**). To assess the impact of temporal information, decoding was tested with clips ranging from 10 to 60 s. Accuracy increased on average with longer clips, demonstrating greater discriminability when more temporal information was included (**Figure 4C**). To test the contribution of voxel selection, decoding was evaluated as progressively larger voxel sets were included. Accuracy declined on average beyond the top-performing voxels, showing that decoding is driven by a subset of the most reliable regions (**Figure 4D**).

Four out of six participants showed robust, well-above-chance decoding, while two showed weaker performance consistent with weaker model fit (**Figure 1D**, **Figure S2D**) and lower cross-session repeatability (**Figure S2**), aligning with prior neuroimaging studies reporting inter-subject variability in semantic mapping [12, 14]. Together, these findings show that DOT signals contain sufficient and robust semantic structure to support model-based decoding, extending optical imaging beyond block design contrasts to naturalistic, high-dimensional tasks.

### Dimensionality reduction and shared semantic group space

Although the voxelwise models estimate model weights for all 1,708 semantic categories, we expect that the underlying semantic representations lie in a shared, lower-dimensional space across individuals (**Figure 5**). To test this, we performed principal component (PC) analysis on bootstrap-derived model coefficients (repeated 50 times) from the top 15% of voxels (**Figure 5A**). To avoid leakage, we selected the best performing voxels per subject from training data through a bootstrapping procedure during model validation (see **Methods**). To compare dimensional structures across representational levels, PCs were derived separately in (1) individual subject spaces, (2) a pooled group space (jackknife approach, leaving the subject of interest out for defining the group space, see **Methods**), and (3) a null space with shuffled stimulus-response alignment (**Figure 5A**).

**Figure 5.**
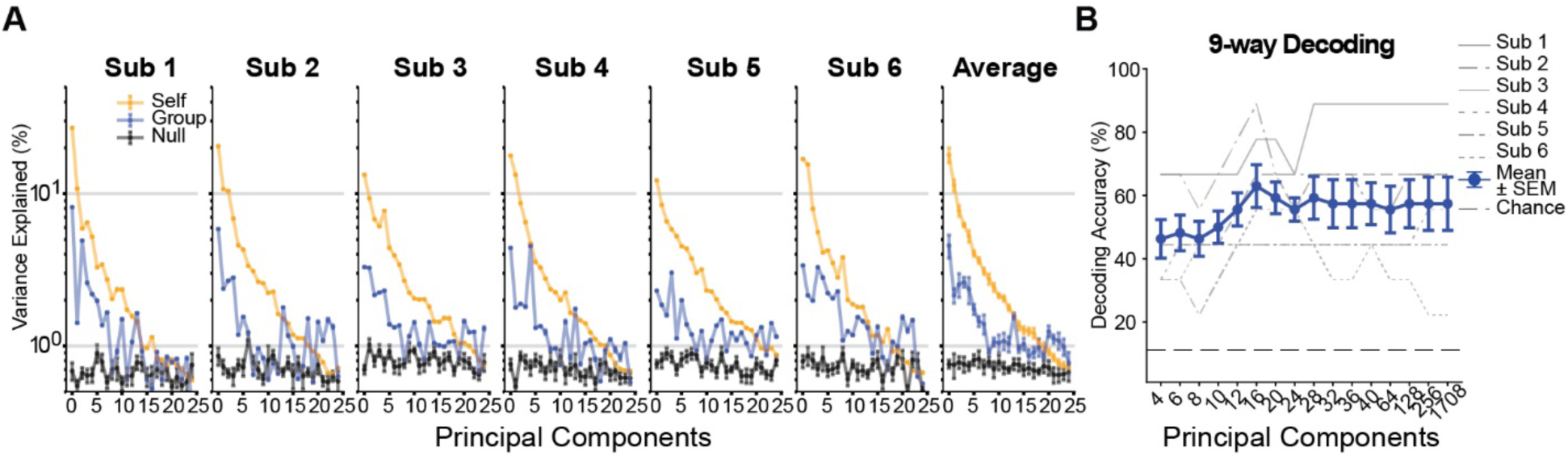
| Principal Component Validation. **(A)** Principal component (PC) analysis was applied to bootstrap-derived model coefficients (repeated 50 times) from the top 15% of voxels (selected by cross-validated accuracy). PCs were computed in three spaces: (i) individual subject spaces (subject-specific coefficients), (ii) a pooled group space (jackknife, leave-one-out approach), and (iii) a null space created by shuffling stimulus features in 60-s blocks to preserve temporal autocorrelation and noise while disrupting stimulus-response alignment. The null space was computed using independent sets of top-performing voxels. This was repeated 10 times. The first 25 PCs are plotted as explained variance (mean ± SEM across subjects). Right panel: group average curves for all three spaces. **(B)** To test whether dimensionality-reduced semantic spaces support decoding, group PCs were derived in a leave-one-subject-out manner from average voxelwise coefficients. WordNet features were projected into these PC spaces, temporally delayed, and fit to each subject’s voxel responses with ridge regression. Clip identification decoding was then performed on the held-out test data using the top voxel subset. The plot shows single subject and mean ± SEM group-level decoding accuracy as a function of the number of retained PCs, with chance indicated by the dashed line. See **Methods** for details on the group space.

To quantify the contribution by each component, we computed variance explained curves for the first 25 PCs. Individual and group spaces showed steep early variance drops, whereas the null space produced comparable flat curves, confirming that variance in true models reflects meaningful semantic structure (**Figure 5A**). Quantitatively, for the group average, the first five PCs accounted for 48.6% ± 2.4% SEM of variance in the individual subject space and 14.2% ± 1.1% SEM in the group space, compared to 3.8% ± 0.1% SEM in the null model. To evaluate the degree of shared representational geometry, we compared group-level PC variance to subject-specific curves. Group PCs captured variance consistently across participants, suggesting a common low-dimensional semantic space.

To test whether dimensionality reduction preserves discriminability, we projected stimuli into the group PC space, re-fit voxelwise models, and performed clip identification decoding using weights from the reduced model (**Figure 5B**, see **Methods**). To determine the effect of dimensionality on decoding, we varied the number of PCs included. Decoding accuracy increased with the first few PCs, peaking around 16 components (62.96% ± 6.76%, mean ± SEM) and remaining stable at 32 components (57.41% ± 7.60%, mean ± SEM), and remaining well above chance (11%, **Figure 5B**). These results demonstrate that most discriminative semantic information lies in a shared subspace aligning with past fMRI studies [12, 14].

### Semantic Dimensions from Group space

Having established a shared low-dimensional semantic space, we next asked whether these components reflect interpretable semantic dimensions [66]. To construct the shared semantic space, we pooled model coefficients across participants and reduced them with PCA, retaining components that explained 80% of the variance (56 PCs). The reduced coefficients were then clustered with k-means into 3 clusters (see **Methods**). To enhance the interpretability of the three cluster features, cluster coefficients were adjusted using WordNet’s hierarchical structure and mapped onto the WordNet semantic tree (**Figure 6**). To support the characterization of the semantic content of each of the three clusters in greater detail, we projected the raw group-derived cluster centroids back into the original WordNet feature space and sorted all 1,708 categories by their signed loadings (**Figure S7**). This analysis provided support for the identified opposing poles within each dimension.

**Figure 6.**
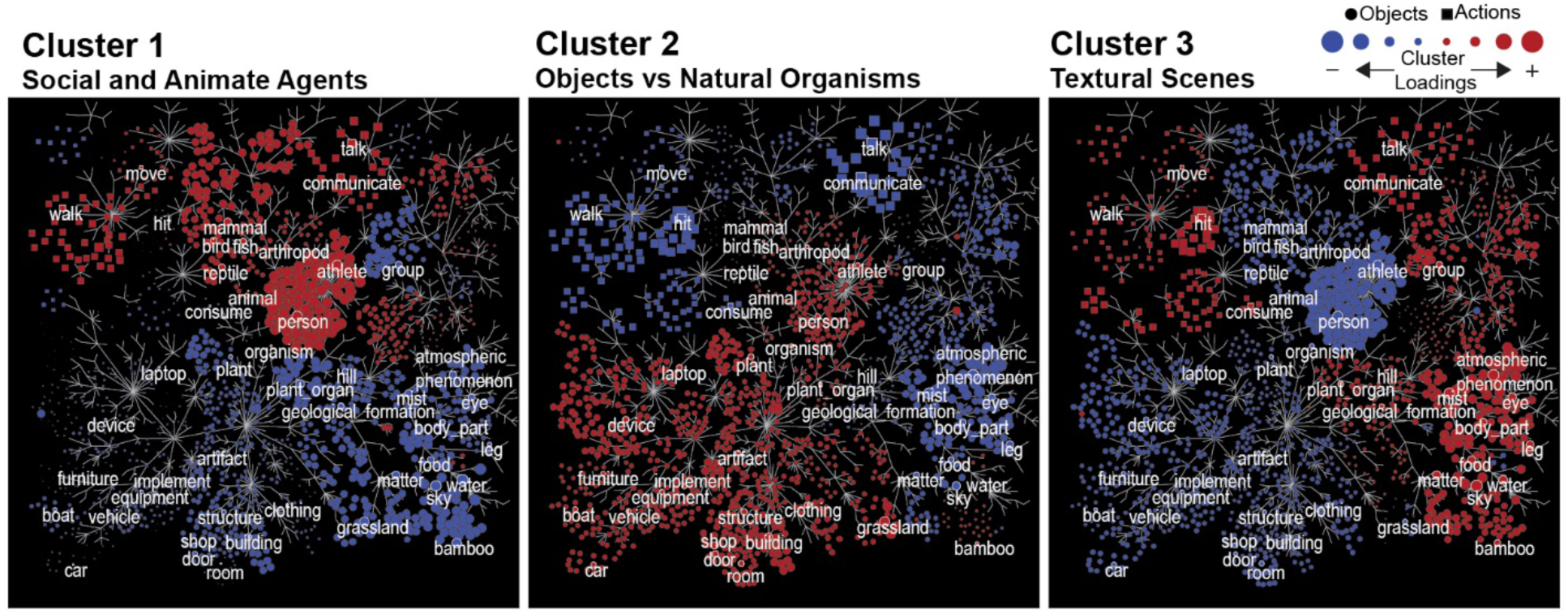
| WordNet-Space Visualization of Group-Derived Semantic Clusters. Each panel displays one of the three semantic clusters derived from group-level *k-means* clustering in PCA-reduced encoding model space. Cluster centers were projected back into the original 1708-dimensional semantic feature space and normalized using a WordNet-based hierarchical smoothing procedure (see **Methods** for details). The resulting smoothed coefficients were z-scored per cluster and plotted onto the WordNet semantic tree. Node color reflects cluster-specific weight (red = positive, blue = negative), and node size encodes the absolute weight magnitude. This visualization shows the distributed semantic structure of each cluster. Cluster 1 emphasizes animate/social agents, Cluster 2 emphasizes objects vs natural organisms, and Cluster 3 emphasizes textural scenes. See **Figure S7** for raw semantic clusters sorted by sign loadings.

The three clusters revealed distinct and interpretable semantic dimensions. Cluster 1 emphasized social and animate agents, with strong positive loadings for people, mammals, and other animals, and negative loadings for artifacts and textural scenes. This dimension aligns with the animacy continuum described in prior fMRI studies [12, 16, 17, 67], supporting the idea that DOT can capture one of the most robust axes of semantic organization. Cluster 2 contrasted objects and cultural artifacts with natural organisms. Positive weights were observed for buildings, devices, and other man-made entities, whereas negative weights emphasized animals and natural phenomena. Similar distinctions have been reported in fMRI studies examining object- versus animal-selective cortex [68, 69], suggesting that DOT recovers a dimension related to the brain’s organization of man-made versus natural categories. Cluster 3 highlighted textural scenes, with positive loadings for sky, water, and atmospheric phenomena, and negative loadings for animate categories. This dimension is consistent with scene-selective representations in ventral visual cortex [65].

When projected back into voxel space, the three semantic clusters revealed reproducible large-scale organization across participants (**Figure 7**). The social and animate agents cluster was expressed in lateral temporal and frontal cortices, consistent with animacy-selective networks previously described in fMRI [12, 16, 17, 67]. The objects versus natural organisms cluster localized to lateral occipital cortex, spanning regions associated with both object-selective lateral occipital cortex (LOC) and scene-selective OPA responses, highlighting a distinction between man-made artifacts and natural categories that has also been observed in fMRI [5, 16, 17, 68, 69]. The textural scenes cluster emphasized sky, water, and atmospheric phenomena at the semantic level, however, when projected to the cortical surface, the spatial maps were more diffuse and less consistent across participants and mapped into the OPA region [5, 65]. Together, these results demonstrate that DOT recovers distributed, high-level semantic dimensions that closely parallel those observed in fMRI.

**Figure 7.**
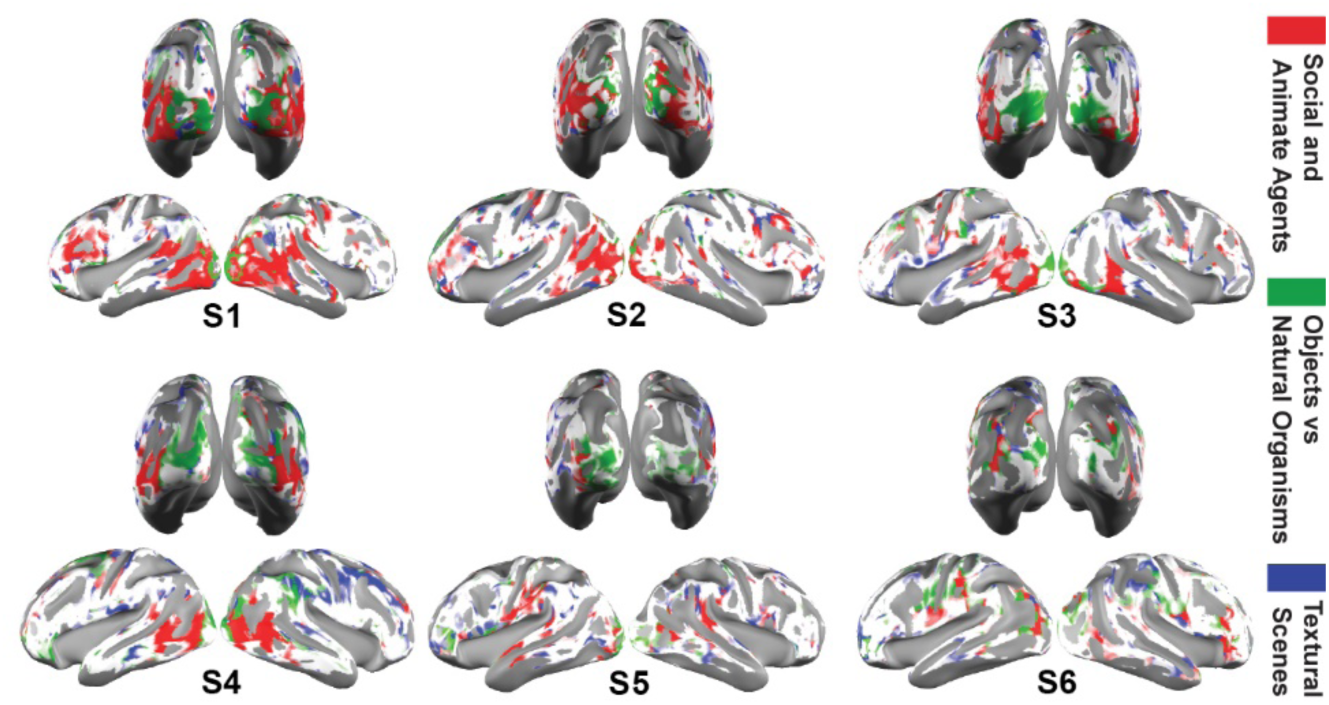
| Semantic Cluster Brain Maps. Semantic clusters assignments, as identified in Figure 6, were mapped to each subject’s cortical surface. Voxel color opacity reflects model generalization performance: voxels with test accuracy ≤ 0 are rendered in white, those between 0 and 0.5 are semi-transparent, and those ≥ 0.5 are shown with full opacity.

## Discussion

Here, we establish a voxelwise semantic encoding approach for non-invasive optical neuroimaging that recovers distributed semantic mappings from naturalistic movie viewings. Specifically, this study has three key contributions: first, we demonstrate the feasibility of high-dimensional semantic encoding using >1,700 features (**Figure 1-2**); second, we establish robust decoding of naturalistic movie clips using a semantic model (**Figure 3-4**); and third, we show a shared semantic space which, combined with dimensionality reduction, organizes into interpretable semantic dimensions (**Figure 5-7**). While similar semantic encoding and decoding results have been shown with fMRI [10–17], they have not been previously shown with fNIRS or DOT. Together, these findings position DOT as a surrogate for fMRI for advanced semantic mapping approaches. By showing that DOT can capture distributed semantic organization in naturalistic paradigms, our work opens the door to extending semantic mapping into populations, environments, and clinical settings that remain inaccessible for fMRI.

### Semantic Encoding with High-Dimensional Features

Our results show that DOT can support voxelwise semantic encoding of a high-dimensional feature space, recovering category-specific responses across lateral occipital, posterior superior temporal, and ventral visual cortices (**Figure 1-2**). These distributed maps align with fMRI findings for key semantic categories, including *person*, *communicate*, *text*, *structure*, and *vehicle*, indicating that DOT can capture semantic selectivity within its cortical FOV. Category-specific responses were reproducible across individuals and model splits, with category weights generalizing above chance at both group and single-subject levels (**Figure S4**). Differences in category strength likely reflect cortical depth. Robust sensitivity is found in superficial regions such as LOTC, pSTG, and lateral ventral cortex, but weaker encoding strength is found in deeper scene/place regions that fall outside optimal DOT sensitivity [5]. Despite the depth-dependent limits of DOT, these findings establish that DOT is sensitive enough to resolve individual semantic categories, positioning it as a viable tool for naturalistic semantic mapping.

### Model-based Semantic Decoding with Naturalistic Stimuli

An essential test of any encoding model is whether the learned representations generalize to new data. We therefore used clip-identification decoding to validate the semantic information captured in the DOT response. If DOT encodes reproducible semantic structure, brain responses should discriminate which movie clips the participant viewed, even across sessions, clip boundaries, and model splits.

Consistent with this assumption, DOT responses supported reliable decoding of movie clips across multiple parametric tunings, including different segmentations, clip lengths, and training set sizes. This suggests that performance reflects stable semantic structure rather than specific data subsets (**Figure 3**, **Figure 4**). Notably, accuracy remained above chance with as little as 10 minutes of training data and even with single test repeats, suggesting the feasibility of decoding in clinical or real-time settings (**Figure 4B**). Decoding performance was strongest when analyses focused on voxel subsets, with informative regions clustering in semantically selective visual areas, such as LOTC, MT, and the ventral cortex (**Figure 4D**). Robust, well-above-chance decoding was observed in four subjects and at the group level, while two participants showed weaker but above-chance decoding across conditions (**Figure S2**). Weaker decoding performance can be explained by individual variability, weaker model fit, or imperfect cross-session alignment.

Extending beyond previously published simple binary-decoding contrasts [43–47], these results demonstrate discriminability across rich, naturalistic stimuli. Prior work has demonstrated the decoding of movie stimuli using HD-DOT with template-based [40, 54, 70] and low-level motion-energy wavelet-based [56] approaches. Here, by leveraging semantic annotations and feature space, we show that DOT signals also preserve higher-order information that supports discriminability across rich naturalistic stimuli.

### Shared Semantic Space and Higher-Order Dimensions

Semantic representations are expected to exhibit a shared structure across individuals rather than being entirely unique. Thus, if the DOT responses capture meaningful semantic structure, individual semantic maps should align in a shared representational space. We therefore pooled semantic coefficient maps into a group space to test whether shared representational geometry could be recovered. The resulting group space preserved discriminative variance well beyond the noise level, confirming a shared representational geometry (**Figure 5A**). Clip identification decoding from this group-derived space remained robust, indicating that semantic discriminability is preserved in a compact, reduced feature space (**Figure 5B**). This is in line with recent fMRI findings that show that naturalistic stimuli can reveal high-dimensional semantic spaces across individuals [12, 14]. By demonstrating similar principles with DOT, our findings show that optical imaging can contribute to this broader shift toward naturalistic, high-dimensional semantic neuroscience.

To interpret the shared semantic space meaningfully, dimensionality reduction and clustering were applied. This approach yielded three robust dimensions that generalized across participants (**Figure 6**, **Figure S7**). The first contrasted animate and social agents with inanimate artifacts and textural scenes, closely aligning with the animacy continuum, which has emerged as a core organizing axis of semantic representation in fMRI [12, 16, 17, 67]. The second dimension distinguished man-made objects and cultural artifacts from natural organisms, consistent with fMRI evidence for object-versus animal-selective cortex [68, 69]. The third emphasized environmental and textural scenes, such as water and sky, versus animate categories, related to scene-related networks in fMRI [65].

Projection of these semantic axes back to the cortex revealed reproducible topographies across occipital, temporal, and frontal cortices that overlap with known semantic networks (**Figure 7**). Spatial maps for the third dimension appear more diffuse and variable across participants, likely reflecting both the distributed nature of scene representations and the limited FOV of DOT. Together, these findings demonstrate that DOT can uncover higher-order semantic axes, animacy, man-made versus natural, and scene-based structure, establishing optical imaging as a viable tool for investigating distributed semantic organization in naturalistic paradigms.

### Clinical and Developmental Relevance

Understanding how semantics are represented in the brain in naturalistic settings is directly relevant to human studies at both ends of the lifespan—developmental and aging—and to a wide range of other clinical applications. One exemplar application is Aphasia, the inability to either understand or produce speech, which occurs in ∼30% stroke survivors [71–73]. Large vessel strokes most commonly (66%) occur in the middle cerebral artery (MCA) and thus most commonly affect the sides of the head, causing damage in some combination of the motor, auditory, Wernicke’s, and Broca’s brain regions. Prior fMRI work shows that visual and auditory semantics share a common neural representation [10, 15, 74, 75]. By bypassing the frequently lesioned language area’s [76], visual stimuli can provide access to semantic representations in participants with aphasia. Similar approaches are valuable for developmental populations [77], including those with autism spectrum disorder (ASD) [78]. Because visual semantics are inherently dynamic, investigating them benefits from paradigms that capture the continuous visual experience during naturalistic movie viewing. Naturalistic movie paradigms also provide a powerful balance between ecological validity, experimental control [8, 9], and enhance engagement [79, 80].

### Implications Across fNIRS and DOT System Architectures

The very high-density DOT system used in this study offers high spatial sampling for naturalistic semantic mapping, but optical systems vary widely in density and coverage. Therefore, an important question is how well the results of this study will generalize to the broader field of optical neuroimaging.

Semantic mapping has been pioneered using fMRI, which typically achieves ∼1.5-2mm isotropic voxel sampling with whole-brain coverage [12, 14, 16, 17], but within a constrained imaging environment [81]. While fNIRS provides an open scan environment, and has been widely used to study language development and recovery [82, 83], aphasia [49, 84], live social interaction [85], and cognition during walking in the real world [86], it is often limited by sparse and localized arrays. As a result, semantic fNIRS studies with sparse channel counts (<50) have primarily relied on block designs and restricted cortical coverage, preventing analysis of distributed semantic structure [43, 45, 47, 87]. Here, we show that DOT can recover distributed semantic organization outside the scanner.

Recent work from our group evaluated the impact of different grid densities on model-based decoding using motion-energy wavelet features with an ultra-high-density DOT system [50]. By subsampling HD-DOT grids, the study found that decoding accuracy had a modest decline from 87% to 74% for template-based decoding and from 62% to 56% for model-based decoding, remaining well above chance (25%) in both cases. These findings suggest that although the present study used a very high-density DOT system for semantic decoding, comparable performance may be achievable with lower-density HD-DOT configurations. While fiber-based HD-DOT has not been widely available, new commercial wearable systems are emerging with high-density array formats [37–39]. It is expected that these wearable formats would capture most of the performance demonstrated in this study.

Prior literature on fNIRS systems using semantic or language stimuli also suggests that the results from this paper may be addressable with different array designs. Sparse-channel fNIRS studies have successfully performed basic decoding of small numbers of semantic objects [43, 44], categories [45], and audio-visual semantic stimuli [46, 47], demonstrating that meaningful semantic information can be recovered even with limited spatial sampling. Similarly, HD-DOT studies have mapped isolated, semantic regressors (e.g., faces, speech, hands) that produced robust cortical maps in both HD-DOT and VHD-DOT systems [40, 54, 55]. Given that decoding in this study relied on top-performing voxels (**Figure 3**), it is plausible that carefully positioned and optimized fNIRS arrays could achieve some level of semantic decoding under naturalistic paradigms with a significantly lower channel count. We are sharing all raw measurement-level data for the imaging presented in this paper, enabling future researchers to subsample arrays from our dataset and systematically evaluate how semantic mapping and decoding scale with grid density and location (see **Resource Availability**).

## Future Directions

Our study demonstrates that DOT can recover distributed semantic organization from naturalistic movies, with implications for both fundamental neuroscience and translational applications. Future work can build on this foundation along three key directions. First, as previously mentioned, systematic evaluation across array densities, spatial configurations, and targeted cortical regions will be critical to inform optimal design trade-offs for semantic mapping and decoding in optical neuroimaging. Second, integrating DOT with complementary technologies, such as eye tracking, could capture how attentional and behavioral states influence semantic representation [88]. Such multimodal integration will become increasingly powerful as advances in fully wearable HD-DOT arrays [37–39, 41, 89, 90] are now making it feasible to expand semantic mapping into everyday environments, creating opportunities to study how semantics are represented during natural social interactions. Finally, full re-synthesis of semantic feature time courses, or category inference, from neuroimaging signals, has been demonstrated with fMRI [11, 13] but not yet with optical imaging. Applied to DOT imaging, re-synthesis techniques would enable real-time semantic decoding and/or closed-loop neurofeedback. Such advances could further increase the impact of DOT as a platform for measuring and understanding semantic deficits with potential future use in monitoring plasticity in clinical populations.

By showing that semantic maps can be obtained outside the scanner, this work broadens the methodological landscape for naturalistic cognitive neuroscience.

## Methods

### Experimental Model and Study Participant Details

#### Participants

Data was collected from six participants (30.8 ± 6 years old, 4 female). All were healthy and had normal to corrected-to-normal vision. Sample size was not determined a priori by statistical power analysis, but was consistent with prior semantic encoding and decoding studies using naturalistic stimuli [11, 12, 16, 21, 91]. Informed consent was obtained from all participants. Consent procedures were conducted in accordance with the IRB protocol approved by the Human Research Protection Office at Washington University School of Medicine. Participants were compensated at a rate of $25 per hour. No data was excluded.

## Method Details

### VHD-DOT Instrumentation

Data were acquired on a custom-built, whole-head Very High-Density Diffuse Optical Tomography (VHD-DOT) system [40] at Washington University in St. Louis, USA (**Figure S1B**, **Figure S1F-G**). The continuous-wave system included 255 source positions (dual wavelength laser sources (685 and 830 nm)) and 252 avalanche photodiode detectors (C12703-112, Hamamatsu) coupled to the head using optic fiber bundles (50-4507-REV1 and 50-4506-REV1, US Fiberoptec Technologies). Fiber bundles were supported by a counterbalanced metal frame and wooden halos, which minimized pressure on the head. The imaging cap provided a first-nearest-neighbor separation of ∼9.75 mm, yielding 9,160 possible measurements with source-detector separations ≤40 mm across both wavelengths (**Figure S1G**). These short-separation channels enabled regression of superficial signals. The system achieved ∼10 mm full-width half-maximum (FWHM) spatial resolution and acquired functional data at 7.8 Hz.

### VHD-DOT Experimental Design

Cap fitting followed established protocols [35, 40] with adjustments for this study. For long-haired participants (N=4), hair was parted at the midline, and four pigtails were placed between motor and side panels. Optodes were combed through hair while monitoring real-time readouts of light level, signal-to-noise ratio, and scalp-coupling coefficients. To align cap placement across sessions, we used a precision tape alignment method [57]. Before cap placement, hypoallergenic silicone tape was applied bilaterally along a line extending from the tragus to the lateral eyebrow. Cap borders were traced on the tape, which remained in place between sessions. For subsequent sessions, the cap was realigned to the tape traces and inion.

Each participant completed three 82 min imaging sessions on the same day. Each session included visual (9 min) and auditory (3 min) localizers, four 10 min training movies, and three 10 min testing movies (**Figure S1A**). Training movies consisted of unique 1 min clips. Testing movies consisted of nine unseen 1 min clips, each repeated ten times in randomized order to minimize adaptation, yielding 90 min across nine 10 min runs. Across sessions, participants contributed 3.5 h of movie data (120 min training, 90 min testing), totaling 21 hours. The imaging sessions included breaks that lasted between ∼30-60 min, depending on the participant’s preference. One participant (S4) completed two sessions consecutively in a single sitting, without a break.

### VHD-DOT stimuli

Natural movie stimuli were identical to those used in prior fMRI [12, 21] and DOT [56] studies. All movies were silent and presented at a ∼15°x15° visual angle. A central fixation square (red, blue, or green, changing at 15 Hz) was presented throughout. Participants were instructed to maintain fixation while attending to the movie content.

Movie frames were annotated with WordNet-based semantic features as described in a prior fMRI study [12]. Each label (e.g., *dog.n.*01) specifies a concept (e.g., *dog*), part of speech (*n* = noun, *v* = verb), and sense index (e.g., *01* = first meaning) (**Figure S1A**). For one minute of test data not included in the existing dataset, labels were generated de novo following the same annotation procedure. See **Feature Extraction** section.

Two functional localizers were also acquired following our standard DOT protocols [35, 40]. The visual localizer consisted of black-and-white checkerboard wedges flickered at 8 Hz for 10 s in the lower left or right visual field, followed by 24 s of rest, repeated over 16 blocks (8 per side) in pseudorandom order. The auditory localizer consisted of passive listening to spoken word lists (1 word/s) during six 15-s blocks, separated by 15-s silent periods.

### MRI data collection and processing

MRI data were collected on a separate day for subject-specific head modeling for DOT. Data were acquired on a Siemens 3T PRISMA Fit scanner using 20- or 64-channel head coils. Each participant underwent T1-weighted MPRAGE (echo time (TE) = 3.13 ms, repetition time (TR) = 2,400 ms, flip angle = 8°, 1 × 1 × 1 mm isotropic voxels) and T2-weighted (TE = 84 ms, flip angle = 120°, 1 × 1 × 1 mm voxels) structural scans, followed by functional localizers (gradient spin-echo EPI; TE = 33 ms, TR = 1,230 ms, flip angle = 63°, 2.4 x 2.4 x 2.4 mm isotropic voxels, multi-band factor = 4). The same visual checkerboard and auditory word-list tasks as in DOT were collected.

Data were preprocessed using fMRIPrep 22.0.2 [92], which is based on Nipype 1.8.5 [93]. To match the VHD-DOT localizer preprocessing, data were detrended and bandpass filtered (0.02-0.2 Hz) before smoothing with an isotropic Gaussian smoothing kernel (10 mm FWHM) [40]. Data were converted to a percent BOLD change measurement by subtracting and dividing by the average BOLD signal in each voxel over time [50]. Full preprocessing details are provided in **Supplemental Methods**.

### VHD-DOT data pre-processing

DOT data were processed similarly to previously reported studies [35, 40, 55]. Raw light levels from two acquisition computers were combined using stimulus sync pulses and converted to differential log-mean intensity values. Channels were rejected if the temporal standard deviation exceeded 7.5% of the mean light level, which indicated contamination by non-physiological variance such as head motion. Rejected channels were excluded from image reconstruction for the entire run. Across participants, 6,917 to 8,598 of the 9,160 possible measurements with source-detector separations ≤ 40 mm were retained. Data were detrended and lowpass filtered with a 1 Hz cutoff. Superficial signal regression was performed using the average of first-nearest-neighbor channels (∼10 mm separation) to reduce scalp and systemic contributions [36, 94]. For the naturalistic movie data, signals were subsequently low-pass filtered at 0.2 Hz and downsampled to 1 Hz for further analysis. Localizer tasks were low- and high-pass filtered at 0.2 Hz and 0.02 Hz, respectively, before downsampling.

### VHD-DOT Image Reconstruction and Spectroscopy

Subject-specific light models were generated for each participant using their anatomical MRI scans following established protocols [40, 55]. Head models were derived from the first imaging session and assumed valid for subsequent sessions under the precision tape alignment procedure. Source and detector arrays were initially registered to an MRI-derived, five-layer segmented head mesh, which is critical for accurate light modeling because it accounts for the distinct optical properties of scalp, skull, cerebrospinal fluid, gray matter, and white matter [35, 94]. Anatomical landmarks (tragus, inion) were aligned to photographs collected during the DOT imaging session. Segmentation was performed using FreeSurfer [95] and NeuroDOT [35, 96], and the mesh was generated with NIRVIEW [97]. Optode positions were then relaxed onto the head using an iterative optimization that balances source-detector distances with placement on the head mesh [98]. Photon diffusion through the mesh was modeled with NIRFAST’s finite-element solver [99]. To further refine registration, functional maps from DOT visual and auditory localizers were compared with subject-specific fMRI localizer responses using overlap maps and dice coefficients. Array placement was refit if necessary. This two-step procedure ensured accurate alignment of optodes to each participant’s anatomy and function. DOT images were reconstructed in voxel space at 2 mm isotropic resolution using the subject-specific light models to solve the DOT inverse problem [35, 58, 98]. Hemodynamic signals were then estimated using spectral decomposition of the absorption changes at 685 and 830 nm to derive relative concentration changes in oxygenated and deoxygenated hemoglobin (HbO, HbR) [51, 56].

### Task Data Analysis

fMRI and DOT localizer data were analyzed with a general linear model (GLM) in voxel space. Task regressors were convolved with a canonical hemodynamic response function previously derived for HD-DOT [53]. Beta values were estimated for each run, representing stimulus-evoked responses relative to baseline rest periods. The resulting beta maps were used for subsequent subject-specific head model and data alignment assessment.

### VHD-DOT Co-registration and Repeatability Analysis

To assess cross-session reproducibility, we evaluated both localizer and movie data. For localizers, mean voxelwise temporal correlations were computed across sessions for visual (checkerboard) and auditory (word list) tasks, and transformed using Fisher’s r-to-z (**Figure S2**).

For naturalistic movies, a subject-specific brain mask including gray matter, white matter, and cerebrospinal fluid was applied to the reconstructed voxel space, yielding 75,151 ± 3,150 voxels retained per subject (mean ± SEM). Data from repeated movie presentations were concatenated across sessions, yielding matrices 𝒀_𝒕𝒆𝒔𝒕_ ∈ ℝ^𝑛𝑅×𝑛𝑇×𝑛7^ and 𝒀_𝒕𝒓𝒂𝒊𝒏_ ∈ ℝ^𝑛𝑇×𝑛7^ , where 𝑛_𝑅_ is the number of repeats, 𝑛_𝑇_ is the number of timepoints and 𝑛_𝑉_ is the number of voxels.

Test movie repeatability was quantified as explained variance (EV), which captures the proportion of signal consistent across repetitions and thus provides an upper bound on encoding performance.

For a voxel 𝑣, with measured signal 𝒚_𝒕𝒆𝒔𝒕𝑣𝑟_ on repetition 𝑖 and mean response 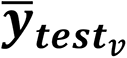, EV was computed as:

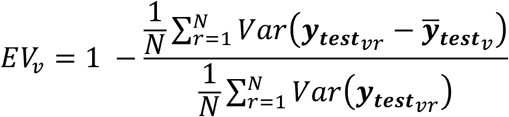

where 𝑁 is the number of repetitions. Here, the numerator corresponds to the residual (within-repeat noise) variance and the denominator to the total voxel variance. Thus, 𝐸𝑉_𝑣_reflects the proportion of reliable, stimulus-driven responses (**Figure S2B**).

### Feature Extraction

Semantic annotations were derived from the WordNet-based framework of [12]. Each second of the movie stimulus was labeled with 1,364 WordNet synsets [59] coded as binary presence/absence (**Figure S1C**). Hierarchical expansion using WordNet hyponym-hypernym relationships added 341 higher-order categories, yielding 1,705 total labels. For one test segment not included in the original dataset, manual annotation identified three additional categories, resulting in a final feature set of 1,708 labels. Following prior semantic encoding work with this stimulus set [12], features were expanded with a finite impulse response (FIR) design. Eight one-second delays (1-8 s at 1 Hz) were applied to capture the hemodynamic response, yielding 13,664 regressors. To reduce dimensionality while preserving temporal structure, adjacent delays were averaged in pairs (1-2 s, 3-4 s, etc.), yielding 4 temporally smoothed regressors per feature (6,832 total).

### Encoding Model

We modeled voxelwise DOT responses to time-varying semantic features of the movie stimuli using L2-regularized linear regression. Brain responses of the training data were represented as a matrix 𝒀_𝒕𝒓𝒂𝒊𝒏_ ∈ ℝ^𝑛𝑇×𝑛𝑉^, where each column corresponds to the HbO time course of a voxel across time points (mean-centered and normalized by standard deviation). Stimulus features were represented as 𝑿_𝒕𝒓𝒂𝒊𝒏_ ∈ ℝ^𝑛𝑇×𝑛𝐹^, 𝑛_𝐹_ is the number of features, with columns corresponding to semantic categories (object/action labels, temporally delayed, and mean-centered). Model weights were estimated in a matrix 𝜷 ∈ ℝ^𝑛𝐹×𝑛𝑉^, capturing how each feature contributes to each voxel’s response. For the forward model, voxelwise predictions were defined as 𝑿_𝒕𝒓𝒂𝒊𝒏_𝜷 = 𝒀_𝒕𝒓𝒂𝒊𝒏_ + 𝝐, with residuals captured in 𝝐. To estimate model weight coefficients, 𝜷 was fit using L2-regularized linear regression, minimizing ‖𝑿_𝒕𝒓𝒂𝒊𝒏_𝜷 − 𝒀_𝒕𝒓𝒂𝒊𝒏_‖^2^ + 𝛼‖𝜷‖^2^. To ensure scale-invariance across feature spaces, the regularization parameter was scaled using the norm of 𝑿_𝒕𝒓𝒂𝒊𝒏_ (**Figure S1D**).

### Hyperparameter tuning

Voxelwise L2-regularized models were fit separately for each participant. Regularization strength (𝛼) was optimized per participant using a bootstrapping-based cross-validation (repeated 100 times). The 120 min training data was first divided into contiguous, non-overlapping 60 s blocks. In each bootstrap iteration, 24 blocks (20% of the data) were randomly selected as validation data, and models were fit on the remaining blocks. Prediction accuracy was assessed as the Pearson correlation between predicted and observed responses in the held-out blocks. Candidate values 𝛼 ∈ [0.1,0.3,0.5,0.7,1.0] were tested (𝛼 was scaled by the norm of 𝑿_𝒕𝒓𝒂𝒊𝒏_). The best-performing 𝛼 was identified per participant, averaged across participants, and rounded to the nearest grid point, yielding a group-level value of 𝛼 = 0.3. 60-second blocks were chosen as they exceed the DOT hemodynamic autocorrelation window while providing sufficient samples per fold.

### Best voxel selection

To identify reliably estimated voxels, models were re-fit on the full 120-min training set using the group-level regularization parameter (𝛼 = 0.3). Within each of 10 repetitions (different random seeds), 30 bootstrap repeats were performed, each with 20% of blocks held out. Voxelwise accuracy was averaged across bootstraps and then across repetitions to yield a final reliability score. To capture variability, the 10 mean accuracy maps from individual repetitions were also retained, producing slightly different voxel subsets for downstream decoding. The top-performing voxels identified by this procedure were used for subsequent decoding, PCA, and Cluster analyses.

### Encoding Model Validation

Model performance was evaluated as the Pearson correlation between predicted and observed voxel time courses.

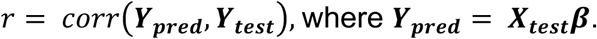

The test set (𝒀_𝒕𝒆𝒔𝒕_ ∈ ℝ^𝑛𝑅×𝑛𝑇×𝑛7^) consisted of nine minutes of unique movie clips not included in training, each repeated 10 times. To improve signal-to-noise ratio, test responses were averaged across repetitions prior to evaluation, yielding 𝒀_𝒕𝒆𝒔𝒕_ ∈ ℝ^𝑛𝑇×𝑛7^ .

Voxelwise significance was assessed using a blockwise permutation test performed separately for each subject. The measured responses were randomly permuted in contiguous 60 s blocks (1,000 iterations) to preserve temporal autocorrelation while disrupting stimulus alignment. For each permutation, voxelwise correlations between predicted and permuted responses were recomputed to form a null distribution. Empirical p-values were obtained as the proportion of permutations where permuted correlations exceeded the observed correlation. P-values were converted to z-scores using Fisher’s r-to-z transform and corrected for multiple comparisons across voxels using the false discovery rate (FDR; p < 0.05).

All model fitting and validation were implemented in Python using NumPy [100], SciPy [101], and scikit-learn [102], with ridge regression routines adapted from publicly available code provided by Huth and colleagues [14] and with utility functions (e.g., stimulus delayers and WordNet visualization) adapted from Gallant Lab’s voxelwise encoding tutorials [103].

### Single Category Encoding

For single-category analyses, voxelwise model weights were first averaged across temporal delays and normalized by their norm. Root mean square (RMS) magnitude was then computed to obtain voxelwise weight magnitudes. For each semantic category, the averaged coefficient was extracted and mapped to the cortical surface. Group-level category maps were derived from fixed-effects t-statistics. For multi-category visualization (five categories), voxels with weak effects (|𝒕| < 0.2 × 𝑚𝑎𝑥|𝒕|) or ambiguous preferences (difference < 0.1 between the top two categories) were rendered white, while remaining voxels were assigned distinct RGB colors based on the winning category. To evaluate robustness, we implemented a split-half identification procedure: models were fit separately to each half of the data, and category weights were cross-validated using a winner-takes-all (WTA) approach applied to the top 10% of voxels (selected by validation performance).

For visualization, group-level category maps were projected onto the MNI152 atlas [104, 105] cortical surface, while individual subject maps were displayed on subject-specific cortical surfaces derived from each participant’s MRI.

### Decoding Model

To quantify the information captured by the semantic category model, we performed a clip identification decoding analysis on the held-out test data. For each test clip, voxelwise brain responses were predicted from the encoding model (𝒀_𝒑𝒓𝒆𝒅_ = 𝑿_𝒕𝒆𝒔𝒕_𝜷). We then constructed a confusion matrix in which each element represented the correlation between the predicted clip and the true clip response 𝑐𝑜𝑟𝑟(𝒀_𝒑𝒓𝒆𝒅_, 𝒀_𝒕𝒆𝒔𝒕_), with rows corresponding to predicted clips and columns to the true clips. The predicted clip (𝑐_𝑝𝑟𝑒𝑑_) was defined using a winner-takes-all strategy as the clip whose predicted response was most correlated with the observed response (𝑐_𝑝𝑟𝑒𝑑_ = 𝑎𝑟𝑔𝑚𝑎𝑥_𝑐_ 𝑐𝑜𝑟𝑟(𝒀_𝒑𝒓𝒆𝒅_, 𝒀_𝒕𝒆𝒔𝒕_)). Correlation was computed across voxels.

Decoding accuracy was defined as the proportion of correctly identified clips, with chance level equal to 1 divided by the number of clips. To improve reliability, decoding was restricted to the top 10% of best-performing voxels (selected by validation performance). The voxel selection procedure was repeated 10 times using different randomly selected subsets of data (see **Best voxel selection**), yielding 10 unique voxel subsets. Accuracy was averaged across these repeats to produce the final measure of decoding performance.

### Decoder Robustness

To examine the strength of DOT movie responses relative to spurious DOT signal in decoding, we varied the number of repeated test clips averaged prior to evaluation (1-10 repeats per subject). For each increment in the number of repeats, 10 random subsets of repetitions were selected without replacement, averaged, and used for decoding. Accuracy was computed as the proportion of correctly identified clips, with chance equal to 1/9.

To evaluate sensitivity to training data size, the total 120 min of training movies were subsampled to 10, 20, 40, 60, 80, or 100 min (1-12 continuous 10 min runs). For each increment of the amount of training data included, a random subset of runs was selected. Temporal structure was preserved within the runs, and 10 random subsets were used for each increment. Models were re-fit on each reduced dataset, and 9-way decoding was performed on the held-out test set.

To test the effect of movie clip length, decoding was performed using test clips ranging from 10 to 60 s (increments of 10 s), while keeping the number of clips in the decoded pool constant (n=9). For each clip length, 10 random temporal segments were sampled from the test set, and accuracy was averaged across repetitions.

To assess task difficulty, the 9 min test sequence was subdivided into 2 to 90 evenly sized clips (270 to 6 s per clip), systematically varying the number of decoding choices. Chance levels were adjusted to 1/n clips.

To evaluate the contribution of voxel subsets on decoding performance, voxels were ranked by bootstrap-cross-validated accuracy in the training set. Decoding was then repeated using subsets ranging from the best performing 100 voxels to all voxels (100, 500, 1k, 2k, 5k, 10k, 20k, 40k, all). Selection was repeated across bootstrap iterations.

All decoding analysis was constrained to the 10% best performing voxels. Voxel selection was based exclusively on training data to prevent test leakage. Group-level decoding accuracy was reported as the mean and standard error of the mean (SEM) across subjects.

### PCA & Cluster Analysis

To identify higher-order semantic structure, analyses were restricted to the top 15% of voxels per subject (selected by validation accuracy during voxel selection). Voxelwise encoding coefficients were mean-centered, L2-normalized, and concatenated across subjects without averaging, following prior semantic encoding work with the same stimulus set [12]. Principal component analysis (PCA) was applied to the pooled feature-by-voxel coefficients, retaining components until 80% of the variance was explained (56 PCs). This criterion balanced dimensionality reduction with variance retention, consistent with prior semantic fMRI studies [66]. *K-means* clustering (with n=3 clusters fixed a priori for interpretability) was then performed in this reduced space to define semantic clusters. For each participant, voxelwise coefficients were normalized and projected into the group PCA space, and cluster assignments were determined based on group-derived *k-means* centroids. Resulting voxel-level cluster labels were mapped to each subject’s cortical surface, with color opacity reflecting model generalization performance (test accuracy ≤ 0 = white; 0-0.5 = semi-transparent; ≥ 0.5 = opaque).

Cluster centroids were projected back into the original 1,708-dimensional semantic feature space, to recover interpretable semantic feature weights per cluster. To capture hierarchical generalizations, we smoothed cluster coefficients by propagating weights upward through the WordNet hierarchy. For each feature, contributions from ancestor categories were included, and each ancestor’s contribution was normalized by the number of its ancestors, preventing features with deeper hierarchies from being overweighted. This procedure was derived from visualization methods used in prior semantic encoding studies [12, 103]. Smoothed cluster coefficients were z-scored and visualized on the WordNet semantic tree, with node color encoding sign (red = positive, blue = negative) and node size encoding absolute weight magnitude.

### PC Validation

Principal component analysis (PCA) was applied to bootstrap-derived encoding model coefficients (repeated 50 times) from the top 15% of voxels selected by validation performance. PCs were computed in three spaces: (i) subject-specific spaces, (ii) a pooled group space with a leave-one-subject-out approach, and (iii) a null space generated by shuffling stimulus features in 60-s blocks to preserve temporal autocorrelation while disrupting stimulus-response alignment. To quantify the proportion of variance explained by PCs, mean-centered voxel coefficients (centered across features for each voxel) were projected onto each unit-normalized PC axis. The squared projection norms across voxels were computed, and these values were divided by the total squared norm of the coefficient matrix. This yielded the proportion of total variance in voxelwise weights attributable to each PC. For the null space, 10 sets of best-performing voxels were defined during model validation, and variance explained curves were computed separately for each set before averaging across repeats. Variance explained curves were averaged across bootstrap samples for subject and group or across null iterations, and uncertainty was quantified as the standard error of the mean (SEM).

To test whether decoding is possible in the reduced semantic space, group PCs were derived in a leave-one-subject-out manner from average voxelwise coefficients. WordNet features were projected into these PC spaces, temporally delayed using the FIR model with four delays, and fit to each subject’s voxel responses with ridge regression. Clip identification decoding was then performed using raw model weights on the held-out test data using the top 10% performing voxels (selected by validation performance).

### Quantification and statistical analysis

Unless otherwise indicated, *n* refers to the number of participants (N=6), and all analyses were performed at the individual-subject level before computing group statistics. Encoding and decoding model performance was quantified using Pearson correlation between predicted and observed voxel responses, clip identification accuracy, as detailed in the figure legends and Results. Voxelwise significance of model performance was assessed using a blockwise permutation test (1,000 iterations, 60 s blocks) to preserve temporal autocorrelation while disrupting stimulus alignment. Empirical p-values were derived from the null distributions of voxelwise correlations and corrected for multiple comparisons using the false discovery rate (FDR; p < 0.05). For PCA analyses, variance explained curves were compared across subject-specific, group, and null spaces. Null models were constructed by shuffling stimulus features in 60 s blocks to preserve temporal autocorrelation while disrupting stimulus-response alignment. For PCA analyses, bootstrap resampling was applied to voxelwise coefficients by sampling voxels with replacement (repeated 50 times) to derive distributions of explained variance curves, whereas for decoding analyses, bootstrapping was used to generate independent voxel masks.

For robustness analyses, bootstrap resampling was used to quantify the stability of voxel selection and model performance. Voxel subsets were defined by cross-validated accuracy in the training set, and decoding analyses were repeated across 10 bootstrap iterations, yielding multiple independent voxel masks. Performance metrics (correlation, EV, decoding accuracy) were averaged across repeats to obtain stable estimates. In all analyses, the center of the distribution is reported as the mean, and dispersion/precision is reported as the standard error of the mean (SEM) across participants or bootstrap repeats, as specified in figure legends.

Fixed-effect t statistics were applied for visualization of single-category maps at the group level (**Figure 2**). All inclusion/exclusion criteria are described in the **Methods**. No participants or data were excluded. Randomization and stratification were applied where indicated, such as random subsampling of training runs and random assignment of test repetitions for averaging.

## Resource Availability

### Lead Contact

Requests for further information and resources should be directed to and will be fulfilled by the lead contact, Joseph P Culver & Wiete Fehner.

## Data and code availability

All data will be deposited are publicly available as of the date of publication. All original code will be deposited and is publicly available as of the date of publication. Any additional information required to reanalyze the data reported in this paper is available from the lead contact upon request.

## Acknowledgements

This work was funded by the Bill and Melinda Gates Foundation [Grant No. OPP1184813 awarded to J. P. C], the National Institutes of Health [grant numbers U01EB027005, R01NS090874, R01EB03491902 awarded to J. P. C., grant number F32DC022178 awarded to J.T., and grant number F31NS110261 awarded to Z.E.M.], the Washinton University’s Imaging Science Pathway Fellowship awarded to W.F. and M.F. [grant number T32EB014855], and the SPIE-Franz Hillenkamp Postdoctoral Fellowship awarded to M.F.. The authors thank all participants for their generous time dedicated to this research.

## Author contributions

Conceptualization, W.F., M.F., J.Ta., Z.E.M., J.Ta., J.P.C., A.G.H.; methodology, W.F., M.F., J.Ta., J.Tr., Z.E.M., J.P.C., A.G.H.; software and resources, W.F., J.Ta., M.F., Z.E.M., A.G.H.; investigation: W.F., M.F., D.W., A.B., A.H.; data curation: W.F., M.F., D.W., J.Ta.: formal analysis, W.F., M.F., J.Ta., J.Ta., J.P.C., A.G.H.; visualization, W.F., M.F., J.Ta., J.Tr.; writing (original draft), W.F.; writing (review and editing), W.F., M.F., J.Ta., D.W., A.B., Z.E.M., A.H., J.Tr., A.G.H., J.P.C.; funding acquisition, J.P.C., A.G.H., M.F., J.Ta., W.F., Z.E.M.; supervision, J.P.C. and A.G.H.

## Supplemental Information

Document_S1.pdf with Figures S1-S7 and Supplemental Methods.

