## Supplemental Information for "Visual Semantic Encoding and Identification of Naturalistic Movies via High-Density Diffuse Optical Tomography"

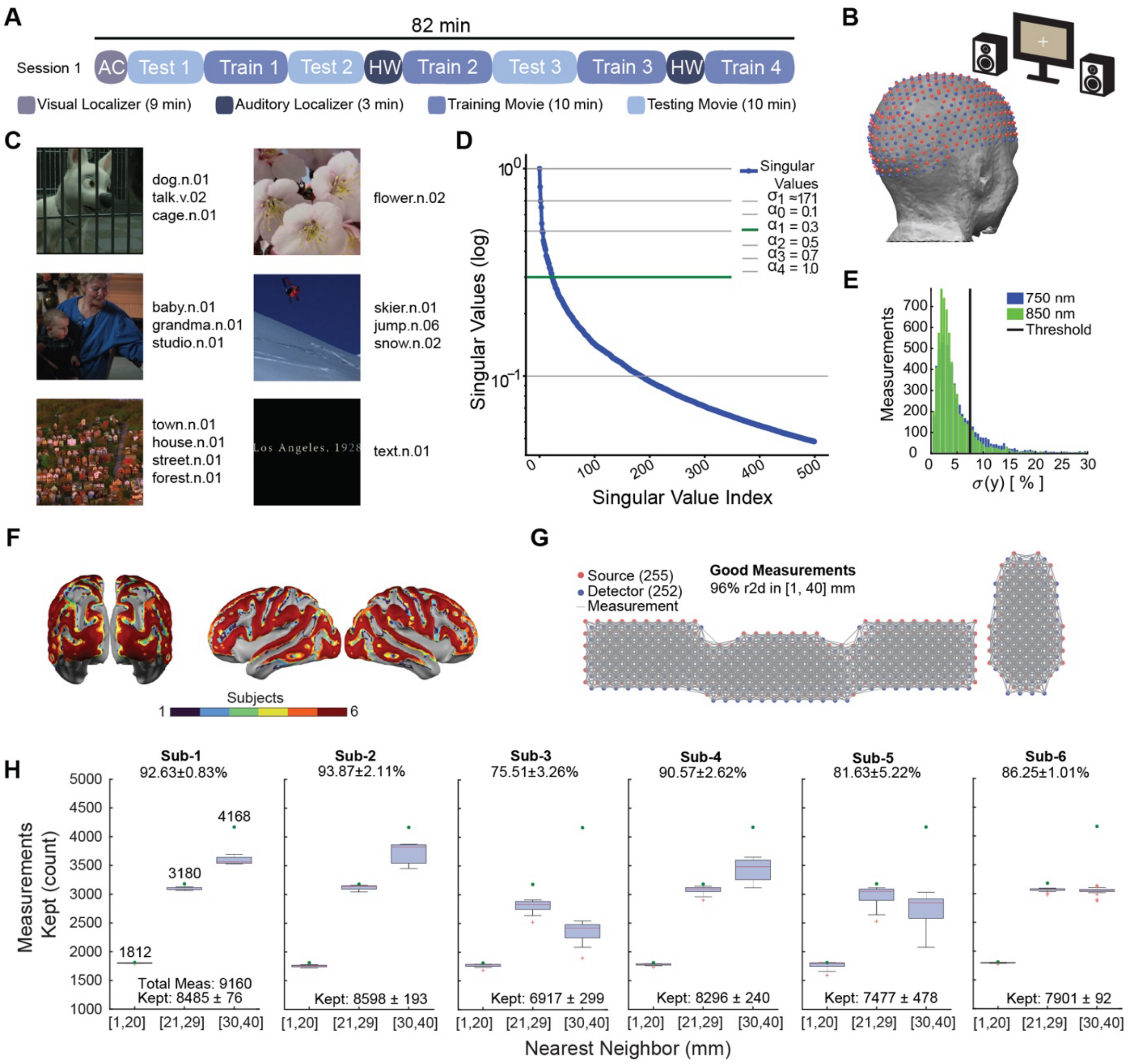


**Figure S1| Experimental Design, Imaging System and Data Quality. (A)** Each participant completed three 82-min imaging sessions, including visual (9 min) and auditory (3 min) localizers, four 10-min training movies, and three 10-min testing movies. Testing movies consisted of 3 unique minutes, segmented into 1-min clips and repeated 10 times per session. Across sessions, this yielded 3.5 h of movie data per participant (120 min training, 90 min testing). **(B)** Source (red) and detector (blue) positions of the high-density DOT cap distributed across the full head. **(C)** Representative movie frames with WordNet-based semantic annotations. Each label specifies a concept (e.g., dog), part of speech (n = noun, v = verb), and sense index (e.g., 01 = first meaning). **(D)** To normalize regularization across feature spaces, parameters were scaled by the norm of the stimulus feature matrix ($\boldsymbol{X}$). The singular value spectrum of $\boldsymbol{X}$ is shown, with the five tested normalized regularization values indicated. The selected hyperparameter (green) lies well below the dominant singular values, indicating that high-variance semantic components were preserved while low-variance/noisy components were effectively suppressed. **(E)** Example histogram of one participant's movie run, showing the distribution of measurement variance. Channels exceeding the 7.5% variance threshold were removed (black line). **(F)** Individual fields of view (FOVs) from subject-specific light models illustrate variability in cap coverage due to head size and shape. Despite this variability, all participants showed full-head coverage with substantial overlap across cortical regions. **(G)** Example "good measurements" plot for one subject, showing sources (red), detectors (blue), and retained measurements (gray). Retained measurements satisfied both the variance threshold (<7.5%) and distance criterion (≤40 mm). **(H)** Measurement retention across all movie-viewing runs for each participant. Boxplots show the number of retained measurements for the first three nearest-neighbor (NN) distances (1-20 mm, 21-29 mm, 30-40 mm). Retention ranged from 76-94% across participants (86.7% ± 2.6%, mean ± SEM), corresponding to 6917 to 8598 retained measurements. Maximum number of possible measurements (green) based on the source-detector separation criterion (≤40mm).

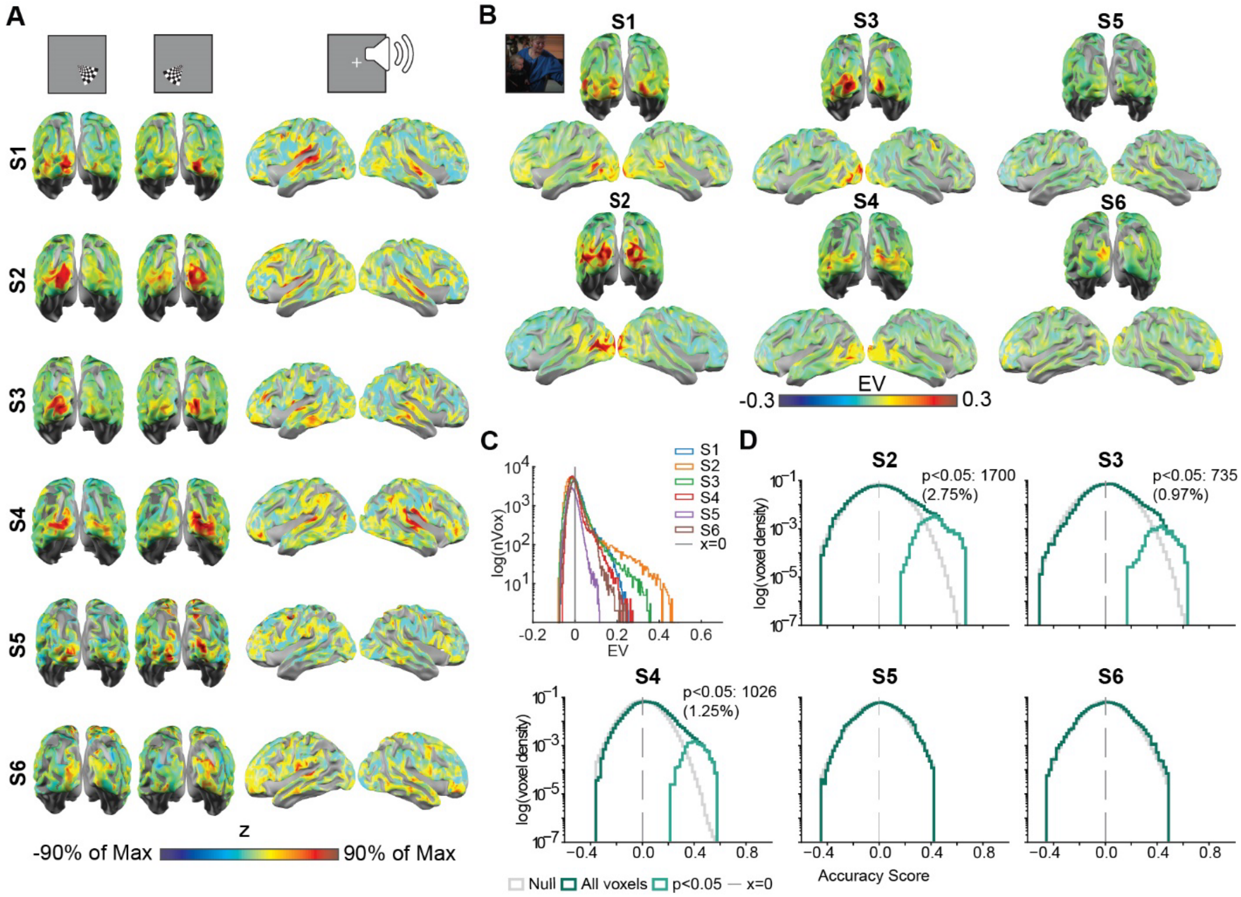

**Figure S2 | Localizer and Movie Repeatability Across Imaging Sessions. (A)** Session-to-session repeatability for visual (checkerboard) and auditory (word list) localizer tasks. Voxelwise temporal correlations were computed across sessions and transformed using Fisher's r-to-z. Subjects S1-4 show strong reproducibility in task-relevant regions, while S5 and S6 exhibit reduced reliability. **(B)** Stimulus-dependent signal content during repeated movie presentations, quantified as explained variance (EV). EV captures the proportion of signal consistent across repetitions and thus provides an upper bound on encoding model performance. Surface maps show high EV in visual cortex. **(C)** Distribution of EV values across voxels for each subject. Histograms highlight intersubject variability in signal repeatability. **(D)** Test accuracy, null distribution, and significant voxels for remaining participants.

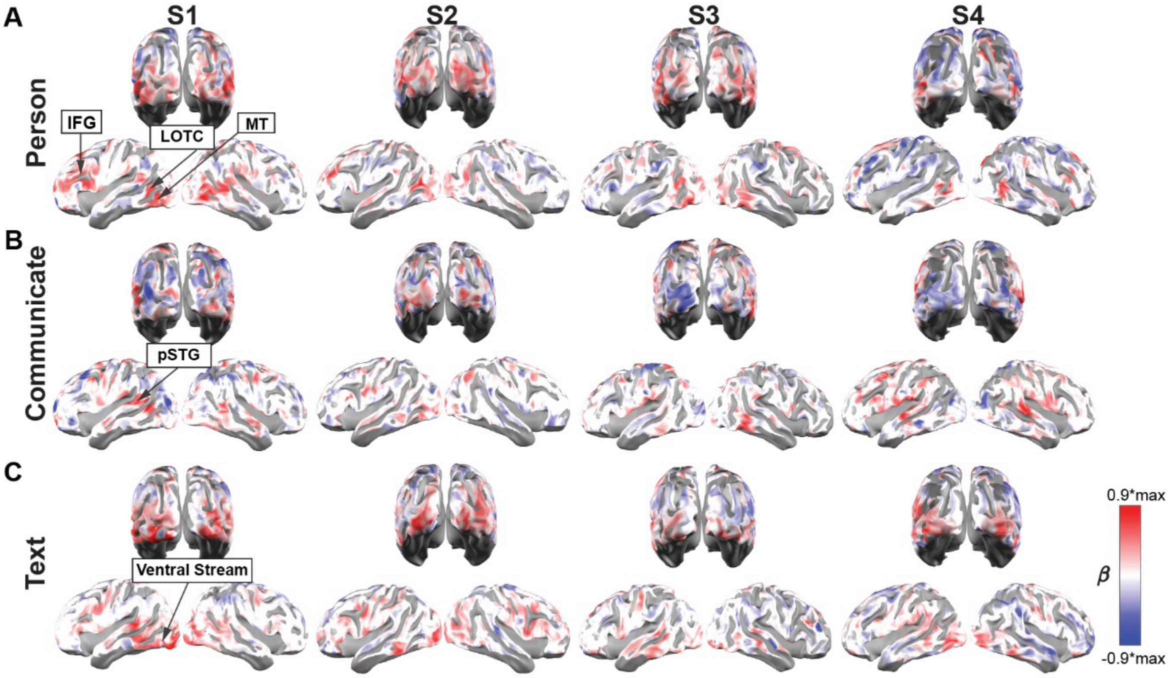


**Figure S3 | Single Semantic Category Mapping, related to Figure 2.** Voxelwise regression weights (β) from the semantic encoding model are shown for three example categories, projected onto subject-specific cortical surfaces for four participants. Warm colors indicate positive weights; cool colors indicate negative weights (scaled to 90% of each map's maximum). **(A)** The category *person* evoked consistent activations across participants in inferior frontal gyrus (IFG), lateral occipitotemporal cortex (LOTC), and middle temporal area MT. **(B)** The category *communicate* showed robust activations in posterior superior temporal gyrus (pSTG), strongest in participants S1 and S4. **(C)** The category *text* was associated with activations along the ventral visual stream.

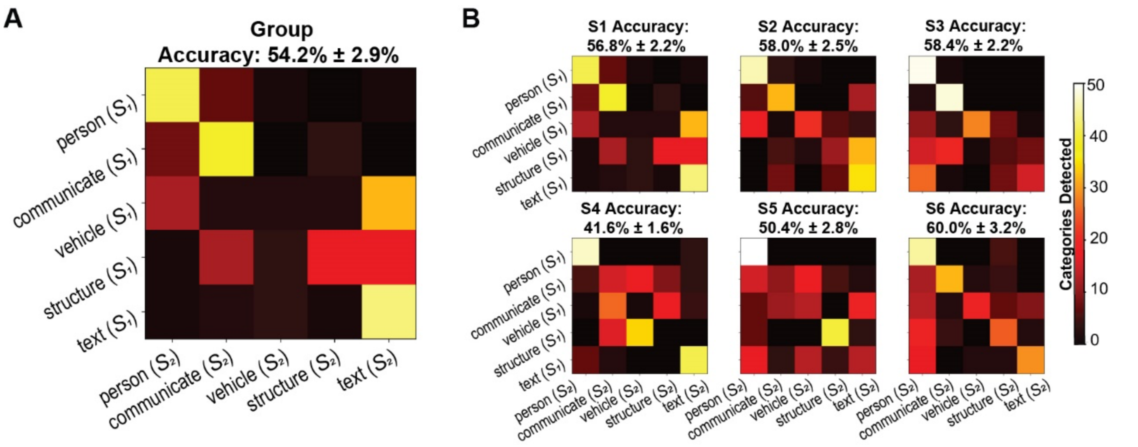


**Figure S4 | Reproducibility of categories weights using encoding model splits, related to Figure 2.** To evaluate the robustness of category representations, we implemented a 50:50 split-half identification procedure. Regression models were trained separately on each half of the data, and category weights were cross-detected between splits using a winner-takes-all approach on the best 10% performing voxels identified during model validation. **(A)** Group-level confusion matrix averaged across subjects, showing above-chance accuracy (54.2% ± 2.9%, mean ± SEM; chance = 20%). **(B)** Individual-subject confusion matrices, all of which exhibited consistent above-chance decoding (range: 41.6-60.0%).

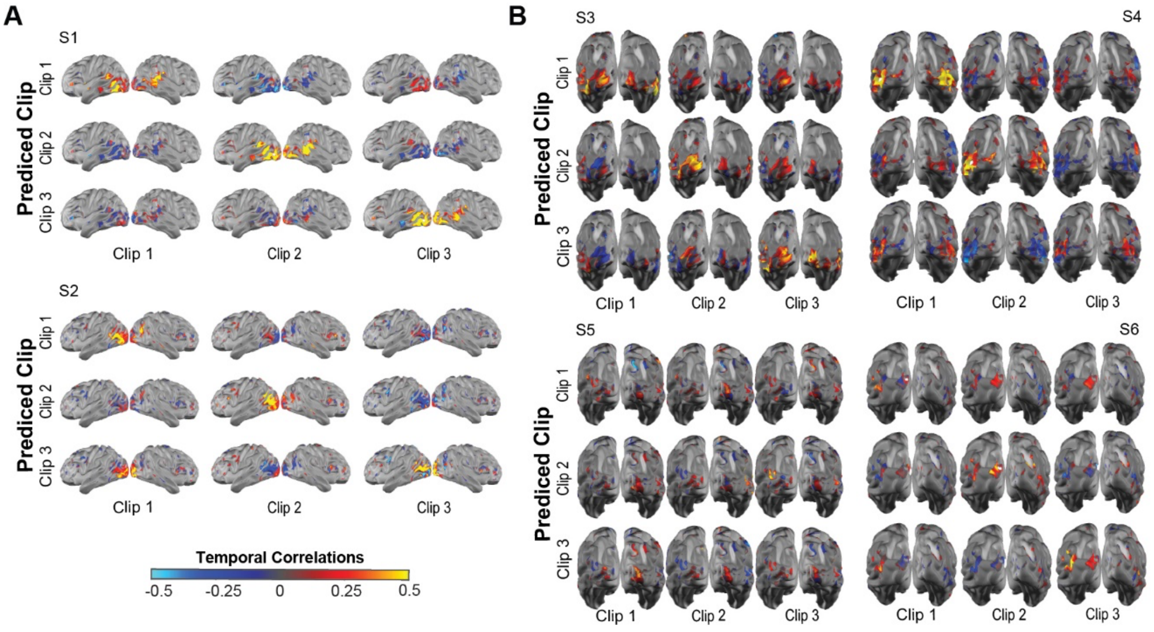

**Figure S5| Single Subject Decoding maps, related to Figure 3. (A)** Lateral views of voxelwise temporal correlations between predicted and measured responses for three 3-min test clips, shown for two subjects (S1-S2). Predicted clips exhibit the strongest correlations with their corresponding measured clips (diagonal structure), indicating successful decoding. Decoding was performed on the top 10% most reliable voxels, selected via bootstrap validation on training data. **(B)** Posterior views for the remaining subjects (S3-S6). S3-S4 show robust clip-specific decoding patterns, while S5-S6 exhibit weaker but still spatially structured correlations.

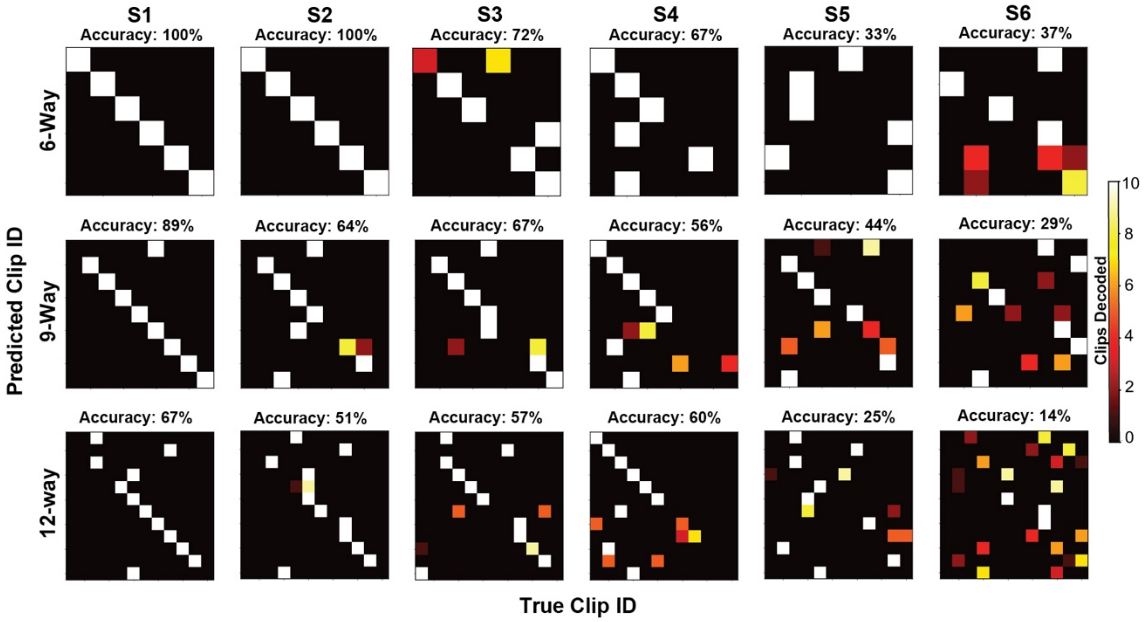


**Figure S6| Single Subject Decoding Performance, related to Figure 3.** Clip decoding at finer segmentations (90 s, 60 s, 30 s) for two subjects aggregated over bootstrapping repetitions. Above-chance diagonal structure is consistent across all granularities. (C) Group decoding accuracy as a function of the number of clips to identify. Nine minutes of test data were segmented into 2-90 clips (270-6 s). Robust above chance decoding was maintained across granularities for all subjects, with S1-S4 showing strongest decoding performance.

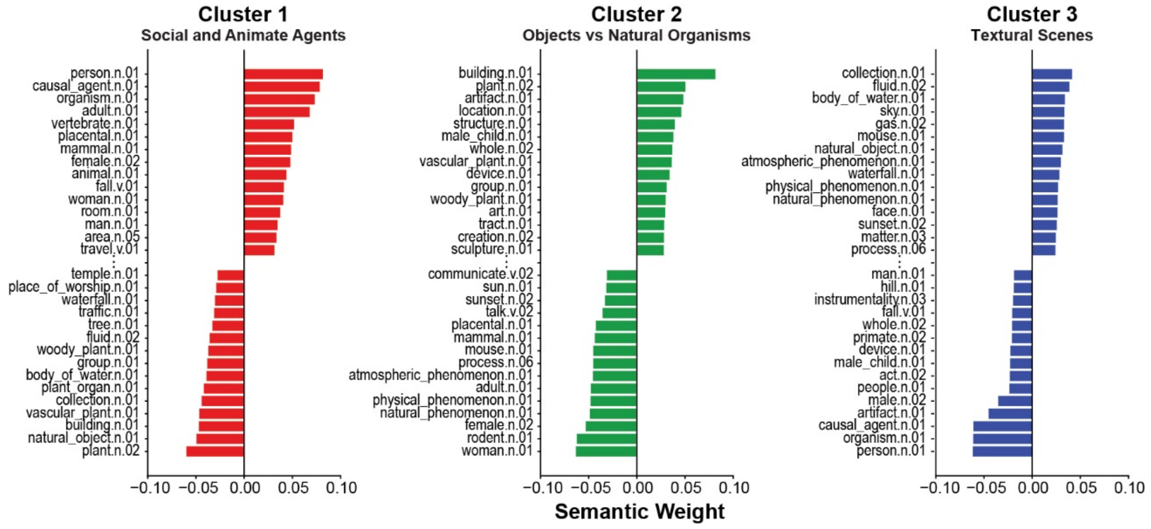


**Figure S7 | Group-Derived Sematic Clusters, related to Figure 6.** Encoding model weights from the top 15% of voxels (per subject) were combined across participants and reduced with principal component analysis (PCA). K-means clustering (k = 3) in the reduced space revealed three interpretable semantic dimensions. Cluster centers were projected back into the original WordNet feature space, and categories were ranked by their signed loadings. For each cluster, the top 15 positive and top 15 negative categories are displayed, with an ellipsis marking the polarity break. Cluster 1 captures a contrast between social/animate agents and the natural world. Cluster 2 reflects a contrast between artificial/constructed objects vs natural organisms. Cluster 3 captures textural scenes.

### Supplemental Methods

Extended fMRI Preprocessing Details (from fMRIPrep)

The following fMIR preprocessing methods are derived from the fMRIPrep boilerplate text as recommended for citing fMRIPrep.

Preprocessing of B0 inhomogeneity mappings: Each subject had at least one field map per imaging session. A B0-nonuniformity map (or fieldmap) was estimated based on two (or more) echo-planar imaging (EPI) references with topup [1]; FSL 6.0.5.1:57b01774).

Anatomical data preprocessing: A total of 1 T1-weighted (T1w) images were found per subject within the input BIDS dataset. The T1-weighted (T1w) image was corrected for intensity non-uniformity (INU) with N4BiasFieldCorrection [2], distributed with ANTs 2.3.3 ([3], RRID:SCR_004757), and used as T1w-reference throughout the workflow. The T1w-reference was then skull-stripped with a Nipype implementation of the antsBrainExtraction.sh workflow (from ANTs), using OASIS30ANTs as target template.

Brain tissue segmentation of cerebrospinal fluid (CSF), white-matter (WM) and gray-matter (GM) was performed on the brain-extracted T1w using fast (FSL 6.0.5.1:57b01774, RRID:SCR_002823, [4]. Brain surfaces were reconstructed using recon-all (FreeSurfer 7.2.0, RRID:SCR_001847, [5], and the brain mask estimated previously was refined with a custom variation of the method to reconcile ANTs-derived and FreeSurfer-derived segmentations of the cortical gray-matter of Mindboggle (RRID:SCR_002438, [6]. Volume-based spatial normalization to one standard space (MNI152NLin2009cAsym) was performed through nonlinear registration with antsRegistration (ANTs 2.3.3), using brain-extracted versions of both T1w reference and the T1w template. The following template was selected for spatial normalization: ICBM 152 Nonlinear Asymmetrical template version 2009c [[7], RRID:SCR_008796; TemplateFlow ID: MNI152NLin2009cAsym].

Functional data preprocessing: For each of the BOLD runs found per subject (across all tasks and sessions), the following preprocessing was performed. First, a reference volume and its skull-stripped version were generated by aligning and averaging 1 single-band references (SBRefs). Head-motion parameters with respect to the BOLD reference (transformation matrices, and six corresponding rotation and translation parameters) are estimated before any spatiotemporal filtering using mcflirt (FSL 6.0.5.1:57b01774, [8]). The estimated fieldmap was then aligned with rigid-registration to the target EPI (echo-planar imaging) reference run. The field coefficients were mapped on to the reference EPI using the transform. BOLD runs were slice-time corrected to 0.566s (0.5 of slice acquisition range 0s-1.13s) using 3dTshift from AFNI [9], RRID:SCR_005927). The BOLD reference was then co-registered to the T1w reference using bbregister (FreeSurfer) which implements boundary-based registration [10]. Co-registration was configured with six degrees of freedom. First, a reference volume and its skull-stripped version were generated using a custom methodology of fMRIPrep. Several confounding time-series were calculated based on the preprocessed BOLD: framewise displacement (FD), DVARS and three region-wise global signals. FD was computed using two formulations following Power (absolute sum of relative motions, [11]) and Jenkinson (relative root mean square displacement between affines, [8]). FD and DVARS are calculated for each functional run, both using their implementations in Nipype (following the definitions by [11]. The three global signals are extracted within the CSF, the WM, and the whole-brain masks. Additionally, a set of physiological regressors were extracted to allow for component-based noise correction (CompCor, [12]). Principal components are estimated after high-pass filtering the preprocessed BOLD time-series (using a discrete cosine filter with 128s cut-off) for the two CompCor variants: temporal (tCompCor) and anatomical (aCompCor). tCompCor components are then calculated from the top 2% variable voxels within the brain mask. For aCompCor, three probabilistic masks (CSF, WM and combined CSF+WM) are generated in anatomical space. The implementation differs from that of Behzadi et al. in that instead of eroding the masks by 2 pixels on BOLD space, a mask of pixels that likely contain a volume fraction of GM is subtracted from the aCompCor masks. This mask is obtained by dilating a GM mask extracted from the FreeSurfer's aseg segmentation, and it ensures components are not extracted from voxels containing a minimal fraction of GM. Finally, these masks are resampled into BOLD space and binarized by thresholding at 0.99 (as in the original implementation). Components are also calculated separately within the WM and CSF masks. For each CompCor decomposition, the k components with the largest singular values are retained, such that the retained components' time series are sufficient to explain 50 percent of variance across the nuisance mask (CSF, WM, combined, or temporal). The remaining components are dropped from consideration. The head-motion estimates calculated in the correction step were also placed within the corresponding confounds file. The confound time series derived from head motion estimates and global signals were expanded with the inclusion of temporal derivatives and quadratic terms for each [13]. Frames that exceeded a threshold of 0.5 mm FD or 1.5 standardized DVARS were annotated as motion outliers. Additional nuisance timeseries are calculated by means of principal components analysis of the signal found within a thin band (crown) of voxels around the edge of the brain, as proposed by [14]. The BOLD time-series were resampled into standard space, generating a preprocessed BOLD run in MNI152NLin2009cAsym space. First, a reference volume and its skull-stripped version were generated using a custom methodology of fMRIPrep. All resamplings can be performed with a single interpolation step by composing all the pertinent transformations (i.e. head-motion transform matrices, susceptibility distortion correction when available, and co-registrations to anatomical and output spaces). Gridded (volumetric) resamplings were performed using antsApplyTransforms (ANTs), configured with Lanczos interpolation to minimize the smoothing effects of other kernels [15]. Non-gridded (surface) resamplings were performed using mri_vol2surf (FreeSurfer).
